# INJURED TUMOR CELLS INACTIVATE ADJACENT CYTOTOXIC T LYMPHOCYTES BY ANTIGEN-INDEPENDENT BLAPTOTROGOCYTOSIS

**DOI:** 10.64898/2026.11.20.768116

**Authors:** Mayura R. Dhamdhere, Quoc Thang Bui, Ågnes Holczbauer, Zhen Lu, Menghao You, Arjun Sengupta, Aalim Weljie, Alice Wang, Kai Tan, Elena Bondarenko, Lang Jiang, Matthew J. Atherton, Yi Fan, Constantinos Koumenis, J. Alan Diehl, Wei Guo, Igor A. Astsaturov, Vladimir S. Spiegelman, Serge Y. Fuchs

**Affiliations:** Department of Biomedical Sciences, University of Pennsylvania, Philadelphia, PA, USA; Department of Systems Pharmacology and Translational Therapeutics, University of Pennsylvania, Philadelphia, PA, USA; Graduate Group in Genomics and Computational Biology, University of Pennsylvania, Philadelphia, PA, USA; Department of Pediatrics, University of Pennsylvania Perelman School of Medicine, Philadelphia, PA, USA; The Marvin C Concetta Greenberg Pancreatic Cancer Institute, Fox Chase Cancer Center, Philadelphia, PA, USA; Department of Clinical Sciences and Advanced Medicine, School of Veterinary Medicine, University of Pennsylvania, Philadelphia, PA 19104, USA; Dept. of Radiation Oncology, University of Pennsylvania, Philadelphia, PA, USA, 19104; Department of Biochemistry, Case Western Reserve University, Cleveland, OH 44106. USA; Department of Biology, University of Pennsylvania, Philadelphia, PA 19104; Department of Pediatrics, Pennsylvania State University College of Medicine, Hershey, PA 17033, USA

**Keywords:** Trogocytosis, Tumor microenvironment, Phosphatidylserine, CART cells, Metabolic dysregulation, Mitochondrial dysfunction

## Abstract

Cancer cells evade cytotoxic T lymphocytes (CTLs) through multiple immunosuppressive mechanisms, including effector trogocytosis, wherein plasma membrane patches carrying tumor antigens are transferred from a donor cancer cell to an antigen-specific CTL across the immune synapse. How this transfer functionally inactivates recipient CTLs remains to be fully understood. Here, we identify blaptotrogocytosis, an antigen-independent form of effector trogocytosis in which membrane fragments are transferred from damaged yet live cancer cells to adjacent, non-specific CTLs. Cancer-cell injury induces phosphatidylserine (PS) externalization, which engages TIM-3 on CTLs to establish cell-cell contacts and drive transfer of PS and other lipids onto the CTL membrane. This pathway also underlies efficient antigen-driven trogocytosis. PS acquisition triggers BLIMP1-dependent metabolic dysregulation and mitochondrial dysfunction, driving CTL exhaustion and impairing antitumor activity of chimeric antigen receptor (CAR) T cells. In pediatric patients with acute lymphoblastic leukemia, elevated levels of TIM-3 and BLIMP1 in pre-infusion products associate with disease relapse. Ablation of BLIMP1 prevents PS-elicited metabolic exhaustion, restores effector function, and enhances antitumor efficacy of human CAR T cells in mouse models. Blaptotrogocytosis thus represents a cell injury-driven mechanism of CTL inactivation and immune evasion.

## Main

CTL-based immunotherapies have achieved remarkable clinical success against B-cell leukemia; however, their efficacy remains attenuated in solid tumors ^1–3^. This limitation is driven in part by immunosuppressive mechanisms within the tumor microenvironment (TME) that promote functional exhaustion of both endogenous CTLs and chimeric antigen receptor (CAR)-bearing CTLs ^2^. Elucidating the mechanisms underlying T-cell suppression within the TME is therefore critical to improve the efficacy of CTL-based immunotherapies.

An important yet under-characterized mechanism of immune evasion is effector trogocytosis - a rapid, contact-dependent process of intercellular membrane transfer between tumor-antigen-expressing cancer cells and antigen-specific CTLs ^4–8^. During this process, CTLs acquire tumor-derived membrane components, including tumor antigens, resulting in the concurrent loss of target antigens from cancer cells ("antigen stripping") that protects them from anti-tumor CTLs ^4,7^. In CAR T-cell therapy, acquisition of targeted tumor antigens also leads to fratricide-mediated killing of therapeutic CTLs ^4^. Finally, transfer of membrane patches onto CTLs leads to their exhaustion ^4,5,7,9^. Despite its physiological and therapeutic significance, the molecular mechanisms governing effector trogocytosis and mediating functional inactivation of recipient CTLs remain poorly defined.

### Blaptotrogocytosis is an injury-induced antigen-independent trogocytosis

We previously demonstrated the importance of 25-hydroxycholesterol in controlling the extent of antigen-dependent effector trogocytosis stimulated by the tumor-derived factors ^10^. We used a model system wherein OVA-expressing cancer cells donated lipids to antigen-specific OT1 CTLs (**Fig 1A**). OVA-deficient donor cancer cells served as a non-specific control. A serendipitous error in depriving these control cells of serum led to a significantly increased ability of cells to donate PKH67 dye-labeled membrane patches to OT-1 cells (**Fig. 1B**).

**Figure 1.**
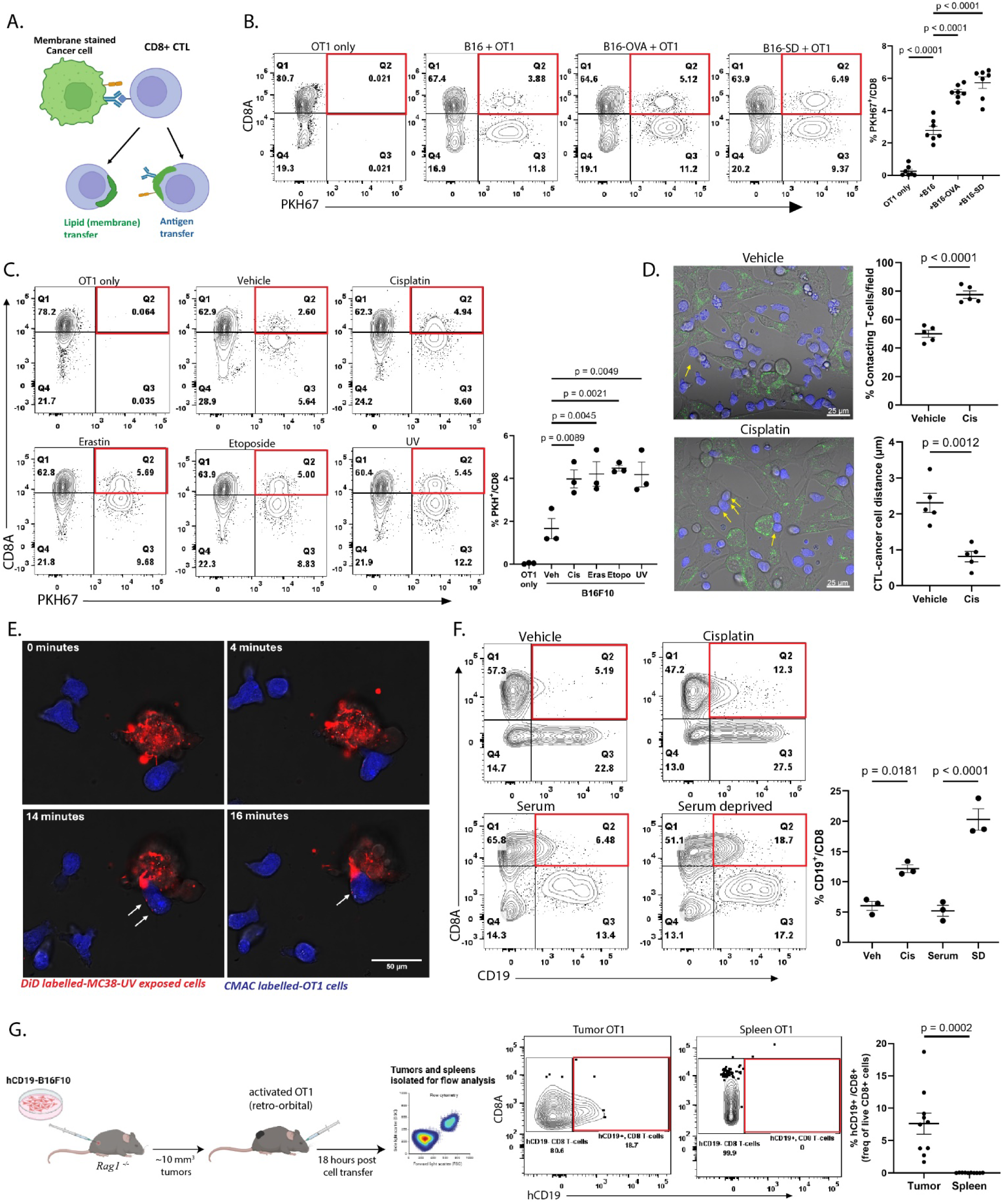
Cancer cell-damaging stimuli induce effector trogocytosis. **a)** Schematic of effector trogocytosis between cancer cells and CTLs. **b)** Transfer of cancer cell membrane onto CTLs, indicated as %PKH67+ CD8+ (of live cells). **c)** Percent PKH67+ CD8+ (of live cells) indicates membrane transfer. **d)** Confocal images of B16F10+OT-I co-cultures showing contacts (arrows) between cancer cells (green) and CTLs (blue), scale bar = 25 µm. Bar graphs show the percentage of CTLs contacting cancer cells per field (top) and cancer cell–CTL distance (bottom). **e)** Time-lapse microscopy of DiD-labelled UV-irradiated MC38(red) + OT-1(blue) co-cultures (arrows indicate membrane transfer). Scale bars = 50 µm. **f)** Percent hCD19+ CD8+ (of live cells) indicates protein transfer. **g)** Experimental layout for *in vivo* trogocytosis (left), and Percent hCD19+ CD8+ (of CD8+ live cells) from tumors or spleens (right, n=10 mice/group). *In vitro* data represents 3-6 independent experiments, and *in vivo* data are representative of 2 separate experiments. Statistical significance determined by one-way ANOVA ***(b-c)***, or unpaired two-tailed t-test ***(d, f-g)*** (mean ± SEM).

Intriguingly, similar results were observed when treating donor cancer cells with cell-damaging stimuli such as UV irradiation, the ferroptosis inducer Erastin, or chemotherapeutics etoposide and cisplatin (**Fig 1C, S1A**). The latter also significantly increased cell-contact frequency and decreased the distance between B16F10 cancer cells and co-cultured OT-1 CTLs within 15 minutes (**Fig 1D**). Furthermore, time-lapse live-cell imaging readily documented the rapid capture of membrane patches by OT-1 T cells from UV-treated MC38 cancer cells (**Fig 1E, S1B** and **Supplementary video**, Supplementary video1 link: https://drive.google.com/file/d/1Yf-UrafFHT6NMPO17q0AIHoiQeqo_HXi/view?usp=sharing).

We termed this injury-induced membrane transfer "blaptotrogocytosis" (from the Greek *Blapto*, βλάπτω - meaning “to harm/damage/injure”). While not dependent on a specific antigen, blaptotrogocytosis exhibited all canonical hallmarks of trogocytosis, including transfer of cell-surface proteins such as human CD19 (hCD19, **Fig 1F** and **S1C**), rapid kinetics (**S1D**), temperature sensitivity (**S1E**), and strict requirements for direct cell–cell contact (**S1F**). Furthermore, known inhibitors of trogocytosis, including the actin polymerization inhibitor Latrunculin (**S1G**) and the membrane-fusion inhibitor 25-hydroxycholesterol (**S1H**), also suppressed blaptotrogocytosis. Importantly, this antigen-independent transfer also occurred *in vivo* following the adoptive transfer of OT-1 CTLs into hCD19-expressing B16F10 (CD19-B16) tumor-bearing *Rag1*-null mice: CD19 protein was observed on OT1 cells infiltrating the tumor but not splenic tissues (**Fig 1G**). Together, these findings reveal blaptotrogocytosis as an antigen-independent form of trogocytosis triggered by tumor cell damage that can occur to a non-specific CTL within the TME.

### Blaptotrogocytosis is driven by TIM-3-dependent recognition of externalized phosphatidylserine (PS)

To determine the molecular foundation of blaptotrogocytosis, we performed an untargeted lipidomics screen on OT-1 CTLs following their brief co-culture with serum-deprived hCD19-expressing B16F10 cancer cells. Trogocytosis-experienced (TRG+) hCD19+ OT-1 cells exhibited a lipidomic profile that was notably distinct from trogocytosis-naïve (TRG-) OT-1 cells **(Fig 2A).** TRG+ CTLs displayed elevated levels of phosphatidylserines (PS), fatty acids, phosphatidic acids, phosphatidylglycerols, oxidized glycerophospholipids, and sphingolipids (**Fig 2A).** We further focused on PS because we observed an increase in multiple species of PS in TRG+ CTLs (**Fig 2B**). Furthermore, PS has been implicated in cellular response to injury ^20^ and in immunosuppression ^12^. In addition, an increase in PS levels on tumor-infiltrating CD8+ T cells was observed after radiation therapy damage ^21^. Finally, a role for PS has been suggested in trogocytosis between CTL and antigen-presenting cells ^22^ as well as in transfer of cholesterol (and potentially other lipids) from cancer-associated fibroblasts (CAFs) onto cancer cells or CTL ^11^.

**Figure 2.**
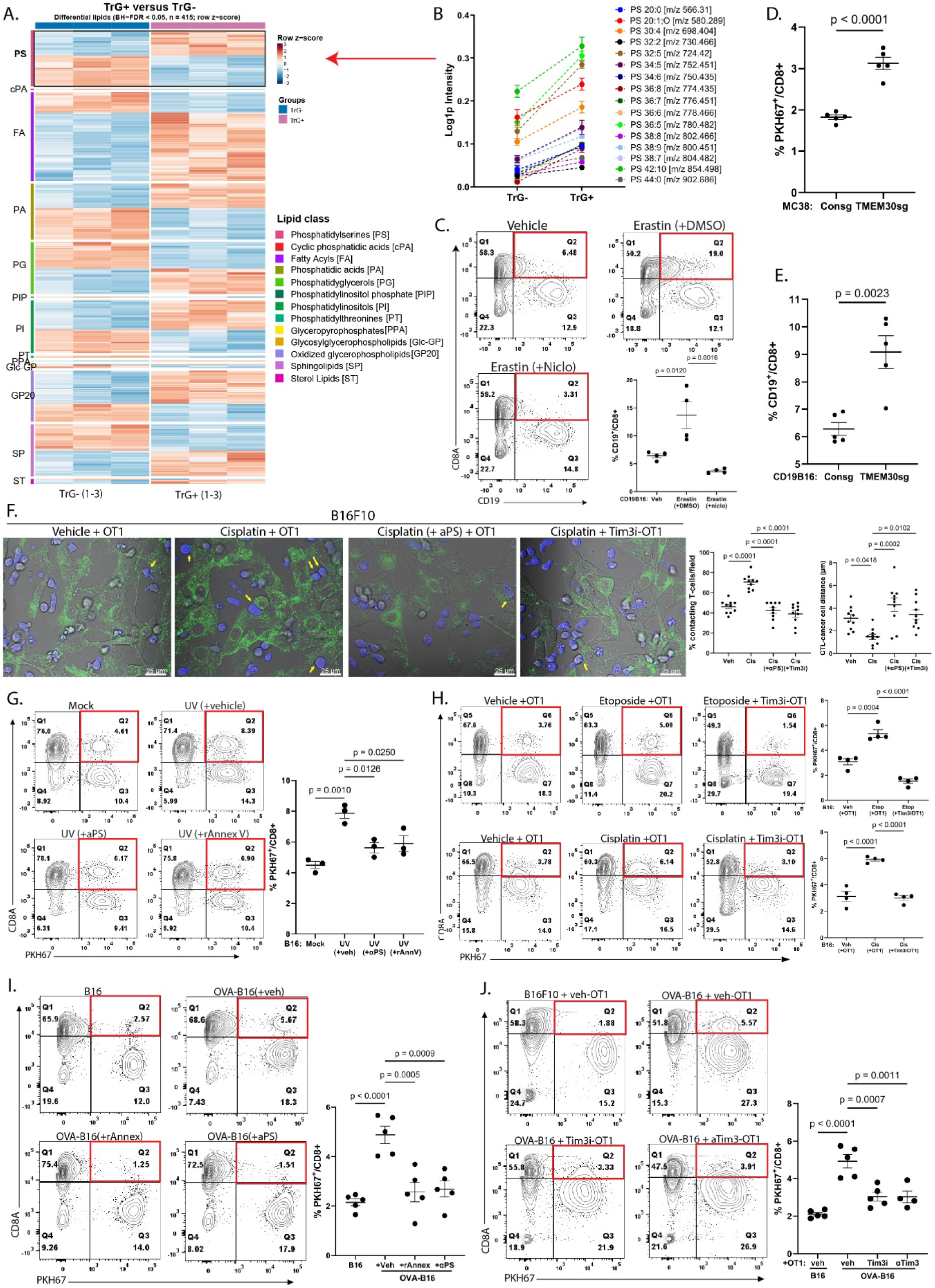
Blaptotrogocytosis and conventional effector trogocytosis are regulated by the PS-Tim3 axis. **a)** Heatmap of differential lipids between TRG- and TRG+ cells (n = 3; one-way ANOVA, BH-FDR < 0.05, Tukey’s post-hoc p < 0.05). Colors show row z-scores from low (blue) to high (red) abundance, lipid families follow standard biochemical abbreviations. **b)** Levels of PS species in TRG- and TRG+ CTLs. ***(c-e)*** Percent TRG+ CD8+ T cells (of live cells) in OT-I co-cultures with: **c)** Erastin ± niclosamide-treated CD19-B16F10 (%CD19+ CD8+); **d)** control or *Tmem30a-KO* MC38 (%PKH67+ CD8+); **e)** control or *Tmem30a*-KO CD19-B16F10 (%CD19+CD8+). **f)** Confocal images of cancer cell–CTL contacts in cisplatin-B16F10 (± αPS) + OT-I CTL (± Tim-3i) co-cultures, quantified as %CTLs in contact with cancer cells (left) and CTL-cancer cell distance (right). **g)** Percent PKH67+ CD8+ (of live cells) in UV-B16F10 (±αPS or rAnnexV) + OT-1 cocultures. ***(h-j)*** Percent PKH67+ CD8+ (of live cells): **h)** in etoposide- or cisplatin-B16F10 + control or Tim3i-treated OT-I cocultures; **i)** in OVA-B16F10 (± αPS, rAnnexV) + OT-1 co-cultures; **j)** in OVA-B16F10 + OT-1 (±TIM-3i or αTIM-3) co-cultures. Data represent 3-5 independent experiments (mean ± SEM). Statistical significance determined by one-way ANOVA ***(c, f, g-j)*** or unpaired two-tailed t-test **(*d-e*)**.

Quantitative ELISA revealed a net increase in total PS levels within native or CAR-bearing CTLs upon their co-culture with serum-deprived or cisplatin-treated cancer cells (**S2A)**. Under homeostatic conditions, PS is restricted to the inner plasma membrane leaflet by TMEM30A-dependent flippases^10,11^. Upon cell damage or stress, PS externalization to the outer leaflet is driven by lipid scramblases such as TMEM16F and XKR8 ^20^. Given that damage-inducing stimuli upregulated externalized PS on live cancer cells (**S2B**), we hypothesized that externalized PS mediates the cell–cell contacts driving blaptotrogocytosis. Indeed, pre-treatment of cancer cells with the scramblase inhibitor niclosamide ^24^ attenuated cisplatin-stimulated PS exposure (**S2C, left**) and significantly suppressed blaptotrogocytosis triggered by either cisplatin (**S2D**) or Erastin (**Fig 2C**).

Conversely, knockout of TMEM30A, which is essential for flippase function ^10,11^, in MC38 or B16F10 cancer cells increased externalized PS (**S2E**) and significantly stimulated blaptotrogocytosis (**Fig 2D-E, S2F**). Furthermore, masking externalized PS on the surface of cancer cells using an anti-PS antibody or recombinant Annexin V (rAnnexV) (**S2C, right**) attenuated cancer cell-CTL contact formation (**Fig 2F**) and suppressed membrane transfer (**Fig 2G**). These results demonstrate that externalized PS on injured cancer cells mediates CTL engagement and subsequent blaptotrogocytosis.

Numerous PS receptors are expressed by diverse cell types ^12–15^. Yet TIM-3 is a major PS receptor expressed on CTLs ^16,17^, and cells from mice lacking the TIM-3-encoding gene *Havcr2* are deficient as trogocytosis recipients^22^. We next investigated whether TIM-3-dependent recognition of externalized PS on injured cancer cells drives blaptotrogocytosis. We pre-treated CTLs with ML-T7, a small-molecule inhibitor of TIM-3 that targets the PS-binding FG–CC’ loop in the IgV domain and thus interferes with interaction with PS ^18^. This treatment significantly reduced cancer cell-CTL contacts (**Fig 2F**) and attenuated membrane transfer induced by serum deprivation (**S1C**), Erastin (**S2G**), cisplatin, or etoposide (**Fig 2H**).

Intriguingly, PS recognized by TIM-3 might also contribute to antigen-driven trogocytosis given that transfer of lipids and hCD19 onto antigen-specific CTLs was attenuated by blocking externalized PS with anti-PS antibody or rAnnexin V (**Fig 2I**). Furthermore, similar effects were observed after inhibiting scramblases with niclosamide (**S2H**) or targeting TIM-3 using either the small-molecule inhibitor ML-T7 or a neutralizing antibody (**Fig 2J**). Moreover, sublethal injury induced by serum deprivation (**S2I, K**) or cisplatin (**S2J**) further increased the extent of antigen-driven trogocytosis onto specific CTLs. These increases were effectively attenuated by niclosamide (**S2J**) or TIM-3 inhibition (**S2K**). Together, these results demonstrate that the PS–TIM-3 axis may regulate both blaptotrogocytosis and conventional antigen-driven trogocytosis and may enable both modes to act in concert during CTL–cancer cell interactions.

### Blaptotrogocytosis induces CTL exhaustion

Although CD8+ T-cells actively utilize lipids to meet bioenergetic demands, excessive uptake of lipids such as cholesterol ^11,19^ or fatty acids may lead to a state of severe exhaustion ^20,21^. Furthermore, oxidized lipids have been shown to inhibit T cell activation and function ^22,23^. Thus, we next examined the effects of blaptotrogocytosis on the functional state of recipient CTLs. Activated OT-1 CTLs co-cultured with *Tmem30a*-deficient hCD19-B16F10 cancer cells displayed significantly reduced expression of IFN-γ and cytotoxic effectors perforin and granzyme B compared to those co-cultured with control cancer cells (**Fig 3A-B**). Similar results were observed in experiments using TMEM30a-deficient MC38 cancer cells (**S3A**). Moreover, co-culture with *Tmem30a*-deficient MC38 cells induced a marked upregulation of the master transcriptional regulator of T-cell exhaustion, TOX, in CTLs (**Fig 3C**).

**Figure 3.**
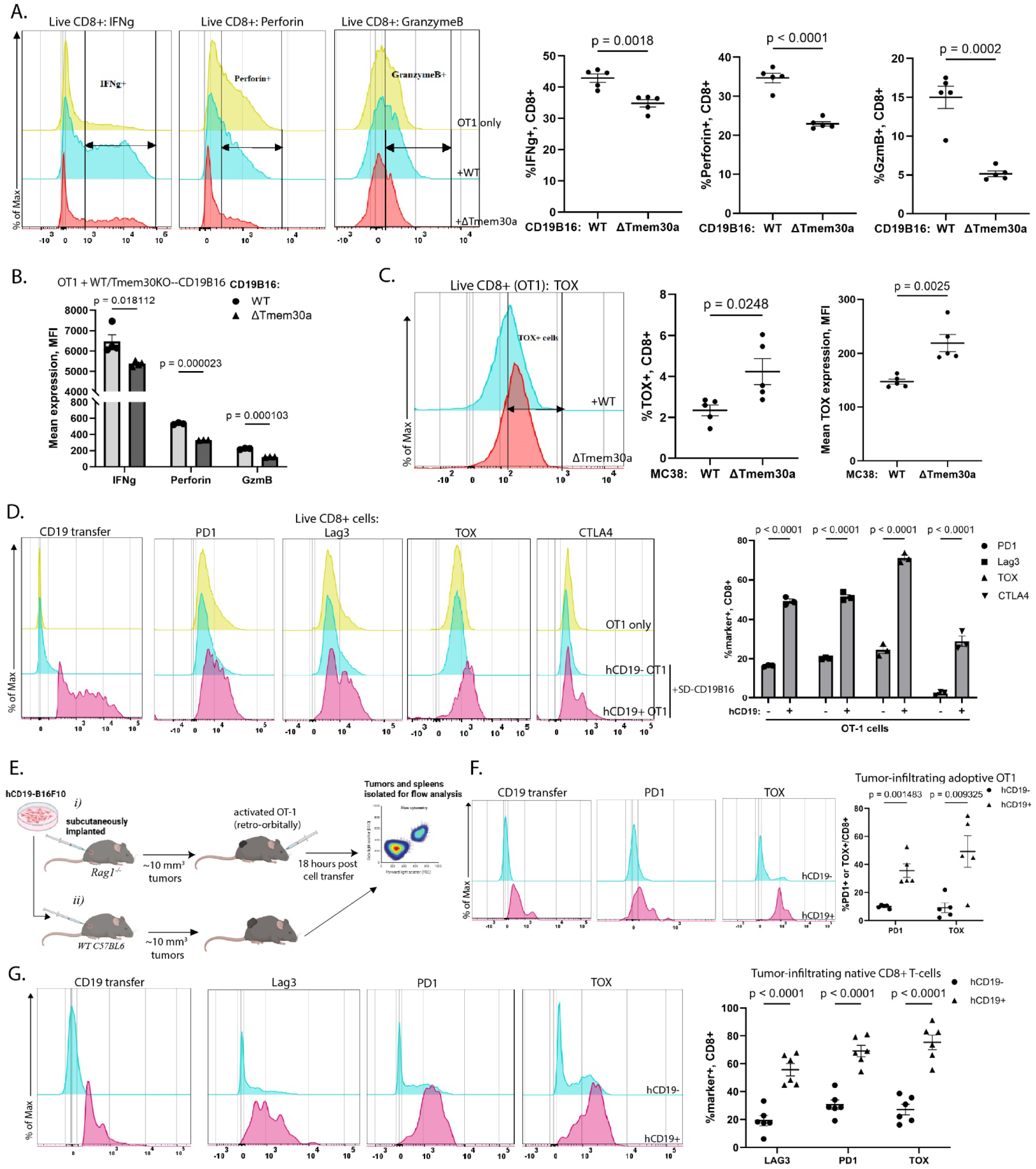
Blaptotrogocytosis inactivates CTLs. **a)** Offset histograms (normalized to mode, % of Max) of IFNg, Perforin and Granzyme B in OT1-cells cultured alone (yellow), with control (blue), or Tmem30a-KO hCD19-B16F10 cells (red), quantified as %marker+ CD8+ (of live cells, normalized to OT-1). **b)** MFI of dataset in ***a)*. c)** TOX expression in OT-1 cells co-cultured with control or Tmem30aKO MC38 cells. **d)** Expression of PD-1, LAG-3, TOX, and CTLA-4 in OT-1 cells alone or from cocultures gated as trogocytosis-experienced (TRG+) or trogocytosis-naive (TRG-). ***(e–g)*** Trogocytosis-mediated CTL exhaustion *in vivo*: **e)** Experimental schema for *(i)* Adoptive transfer model for data shown in ***f***. and *(ii)* endogenous CTL model in hCD19-B16F10 tumor-bearing C57Bl6 mice, for data shown in ***g***. **f)** Expression of PD-1 and TOX in tumor-infiltrating TRG+ (hCD19+, magenta) versus TRG-(hCD19-, blue) OT-1 CTLs (n = 5). **g)** Expression of Lag3, PD1 and TOX in the TRG+ or TRG-tumor-infiltrating endogenous CTLs (n = 6). Both *f) and g)* are quantified as % marker+ cells within each live CD8+ hCD19+ or CD8+ hCD19-gate. *In vitro* data represents 4-5 independent experiments (mean ± SEM). Statistical significance determined by unpaired two-tailed t-test ***(a, c)*** or two-way ANOVA ***(b, d, f, g)***.

Furthermore, inhibition of Erastin- or UV - induced blaptotrogocytosis using Latrunculin, notably decreased the expression of PD-1 in bulk CD8+ CTLs (**S3B**). Of note, these results are likely to be underappreciated because only a fraction of CTLs undergoes blaptotrogocytosis at any given time (**Fig 1**). To overcome this problem, we co-cultured OT-1 CTLs with serum-deprived CD19-expressing cancer cells and then compared the expression of exhaustion markers in CTLs that underwent blaptotrogocytosis (CD19+) versus those that did not (CD19-). Blaptotrogocytosis-experienced (CD19+) CTLs exhibited a robust increase in exhaustion markers, including TOX, PD-1, LAG-3, and CTLA-4, relative to their CD19-counterparts (**Fig 3D, S3C**). Similarly, CTLs that acquired PKH67-stained membrane patches (PKH67+) following co-culture with cancer cells treated with UV, erastin, etoposide, or cisplatin consistently upregulated PD-1 and LAG-3 compared to PKH67-CTLs (**S3D-F**).

Furthermore, we analyzed the adoptively transferred OT-1 CTLs and endogenous intratumoral CTLs isolated from hCD19B16 tumor-bearing *Rag1^-/-^* or wild-type mice, respectively (**Fig 3E**). Blaptotrogocytosis-experienced CD19+ OT-1 CTLs exhibited significantly higher levels of PD1 and TOX compared to CD19-CTLs (**Fig 3F, S3G**). Likewise, in immunocompetent wild-type mice where antigen-driven conventional trogocytosis and stress-induced blaptotrogocytosis co-exist, higher levels of PD1, TOX, and LAG3 were observed in endogenous CD19+ tumor-infiltrating CTLs compared to CD19-TILs (**Fig 3G, S3H**). Finally, in an adoptive transfer model into *Tmem30a*-deficient tumors, CD19+ OT-1 CTLs isolated from *Tmem30a*-knockout tumors displayed further elevated PD-1 and TOX expression relative to CD19+ OT-I CTLs from control (wild-type) tumors (**S3I**). Collectively, these in vivo findings demonstrate that blaptotrogocytosis may inactivate CTLs within the tumor microenvironment.

### Blaptotrogocytosis-mediated PS transfer impairs CTL function

Upon chronic antigen exposure or tumor-directed radiation therapy, exhausted CD8+ T-cells exhibited sustained intrinsic PS externalization ^24,25^. Given that PS levels are increased in blaptotrogocytosis-recipient CTLs (**Fig 2A, S2A**) and that PS was implicated in inhibition of TCR signaling driven by extracellular vesicles ^26^, we examined whether PS transfer occurs during blaptotrogocytosis and contributes to CTL inactivation. We detected increased externalized PS in antigen-driven trogocytosis-experienced (CD19+) CAR T cells upon co-culture with CD19-B16F10 cancer cells (Fig 4A), as well as on PKH67+ OT-1 cells after incubation with PKH67-labeled OVA-B16 cells (**S4A**). Likewise, blaptotrogocytosis-experienced CD19+ OT-1 cells co-cultured with serum-deprived (**Fig 4B**) or cisplatin-treated (**S4B**) cancer cells showed significantly higher levels of cell-surface PS than non-recipient CD19-CTLs. Similarly, increased PS was observed in OT-1 CTLs co-cultured with TMEM30A-deficient cancer cells relative to those co-cultured with control cancer cells in vitro (**S4C**). Importantly, in vivo analysis of CD19-B16 tumor-infiltrating adoptively transferred OT1 (**Fig 4C**) or endogenous (**Fig 4D**) CTLs revealed that PS is markedly increased on blaptotrogocytosis-experienced CD19+ T-cells compared to CD19-counterparts in vivo.

**Figure 4.**
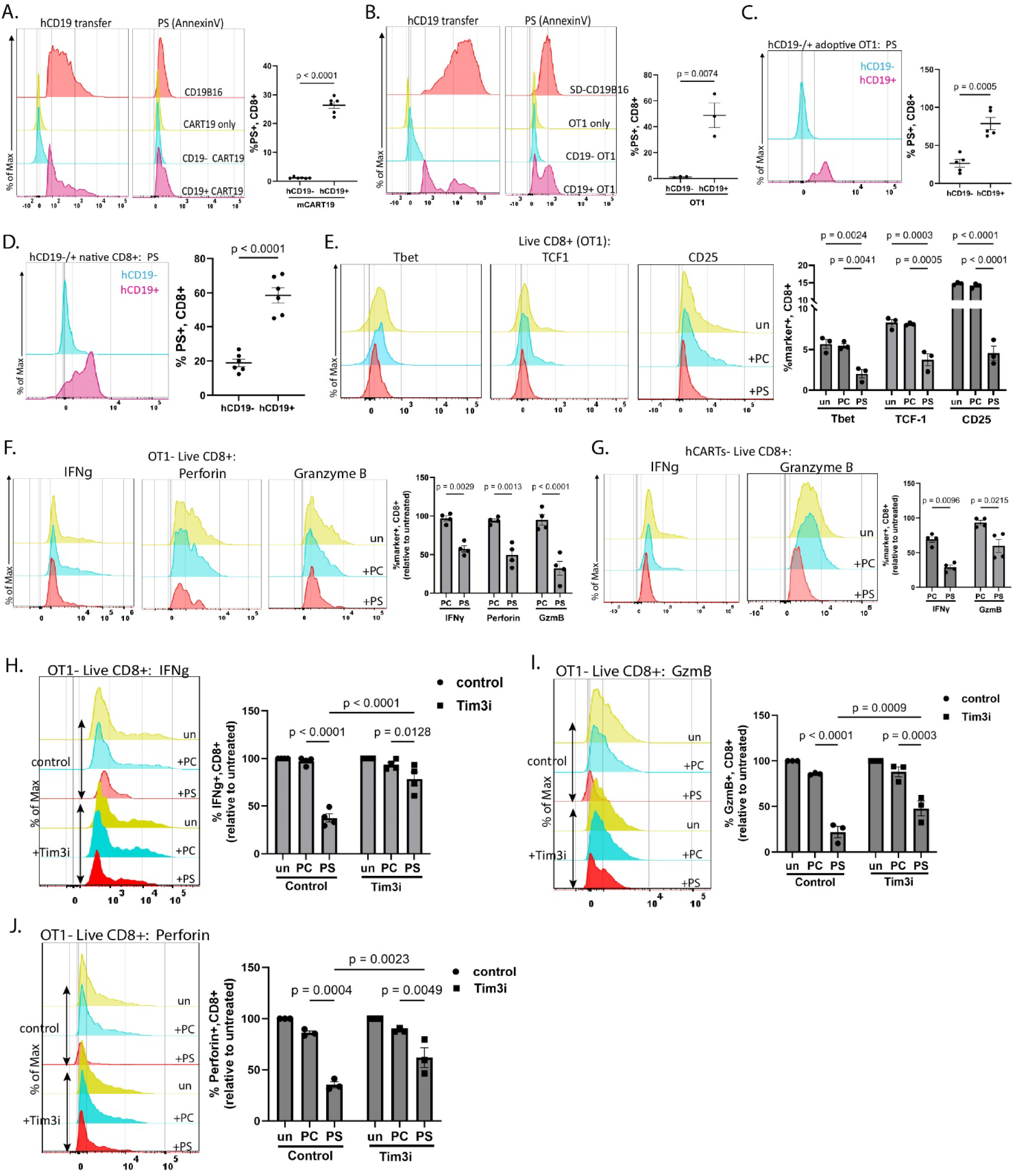
Blaptotrogocytosis involves transfer of PS onto CTLs. ***(a-d)*** Surface PS on TRG+ or TRG-CTLs from: **a)** hCD19-B16F10 + mCART19 co-cultures; **b)** SD-hCD19-B16F10 + OT-1 co-cultures; **c)** tumors of OT-I adoptively transferred mice (n = 5); or **d)** tumors of immunocompetent hCD19-B16F10-bearing mice (n = 6). **e)** Validation of PS-mediated CTL inactivation using liposomes: expression of T-bet, TCF-1 and CD25 in untreated (un) or PC/PS liposome-treated OT-1, quantified as %marker+ CD8+ (of live cells). ***(f-g)*** Expression of IFN-g, perforin, and granzyme B in PC/PS liposome-treated: **f)** OT-1 CTLs, or **g)** human CART cells, quantified as %marker+ CD8+ (of live cells, normalized to untreated controls). ***(h-j)*** Expression of **h)** IFN-g, **i)** Granzyme B, and **j)** Perforin in control or Tim3i-pretreated OT-1 CTLs following PC/PS liposome treatment, quantified as %marker+ (of live CD8+ cells, normalized to respective untreated controls). Data represents 3-4 independent experiments (mean ± SEM). Statistical significance determined by two-tailed unpaired t-test ***(a-d)*** or two-way ANOVA ***(e-j)***.

Given the established immunosuppressive properties of externalized PS ^12,26^ we sought to examine whether, in addition to enabling cancer cell-CTL contact, PS may also elicit inactivation of CTLs. To isolate the specific effects of PS from those of the complex mixture of other biomolecules being transferred during blaptotrogocytosis, we exposed CTLs to synthetic liposomes containing 100% PS or to control phosphatidylcholine (PC) liposomes. Treatment with PS liposomes upregulated PS levels in CTLs in a TIM-3-dependent manner across multiple CTL models, including human CAR T (**S4D**), mouse CAR T (**S4E**), and OT-1 (**S4F**) cells. We confirmed this PS acquisition by quantitative analysis of total cellular PS (**S4G**). Importantly, activated OT-1 CTLs treated with PS liposomes downregulated the activation marker CD25 as well as key lineage-defining transcription factors like T-bet and TCF-1 (**Fig 4E**), while upregulating Eomes – a marker of CTL transition towards terminally exhausted state (**S4H**) and the exhaustion master regulator TOX (**S4I**). Furthermore, PS liposome treatment attenuated the expression of IFN-γ and cytotoxic molecules perforin and granzyme B in activated mouse OT-1 and human CART CTLs (**Fig 4F-G, S4J**). Accordingly, OT-1 CTL treated with PS liposomes displayed a diminished cytotoxic capacity in killing assays against OVA-expressing MC38 or B16 cancer cells (**S4K**).

Because TIM-3-dependent PS transfer to CTLs alone sufficed to inactivate them, we examined whether blocking TIM-3 could rescue CTL function. Pretreatment of OT-1 cells with TIM-3 inhibitor attenuated PS liposome-elicited induction of TOX (**S4I**) and downregulation of IFN-γ (**Fig 4H**), granzyme B (**Fig 4I**), or perforin (**Fig 4J**). Similar rescue effects were observed in other CTL types, including human (**S4L**) and mouse (**S4M-O**) CAR T cells. Together, these data suggest that CTLs acquire surface PS molecules from damaged cancer cells during blaptotrogocytosis, and that this transferred PS can directly drive functional inactivation of recipient CTLs.

### PS inactivates CTLs via BLIMP1-dependent mitochondrial dysfunction

To examine the mechanisms underlying PS-mediated CTL inactivation, we performed RNA sequencing (RNA-seq) on CTLs treated with control-PC or PS liposomes. Gene Set Enrichment Analysis (GSEA) using the MSigDB Hallmark collection revealed a significant PS-driven downregulation of patterns associated with early metabolic priming (MYC targets V1/V2), cell-cycle entry (E2F targets), mitotic progression (G2M checkpoint), and DNA repair pathways (**Fig. 5A**). Importantly, treatment with PS significantly downregulated core bioenergetic pathways, including mTORC1 signaling, glycolysis, oxidative phosphorylation (OXPHOS), and fatty acid metabolism (**Fig. 5A**, **S5A**). The coordinated suppression of OXPHOS and Electron transport chain machinery might have reflected a structural mitochondrial collapse, manifested by significant downregulation of core respiratory complex components (*Ndufs4*, *NdufaS*, *Ndufb8*, *Sdha*, *Sdhb*, *Cox4i1*, *Cox5a*, *Atp5po*), and complex assembly factors (*Coa3*, *Cox20*) (**Fig. 5B, S5B-C**).

**Figure 5.**
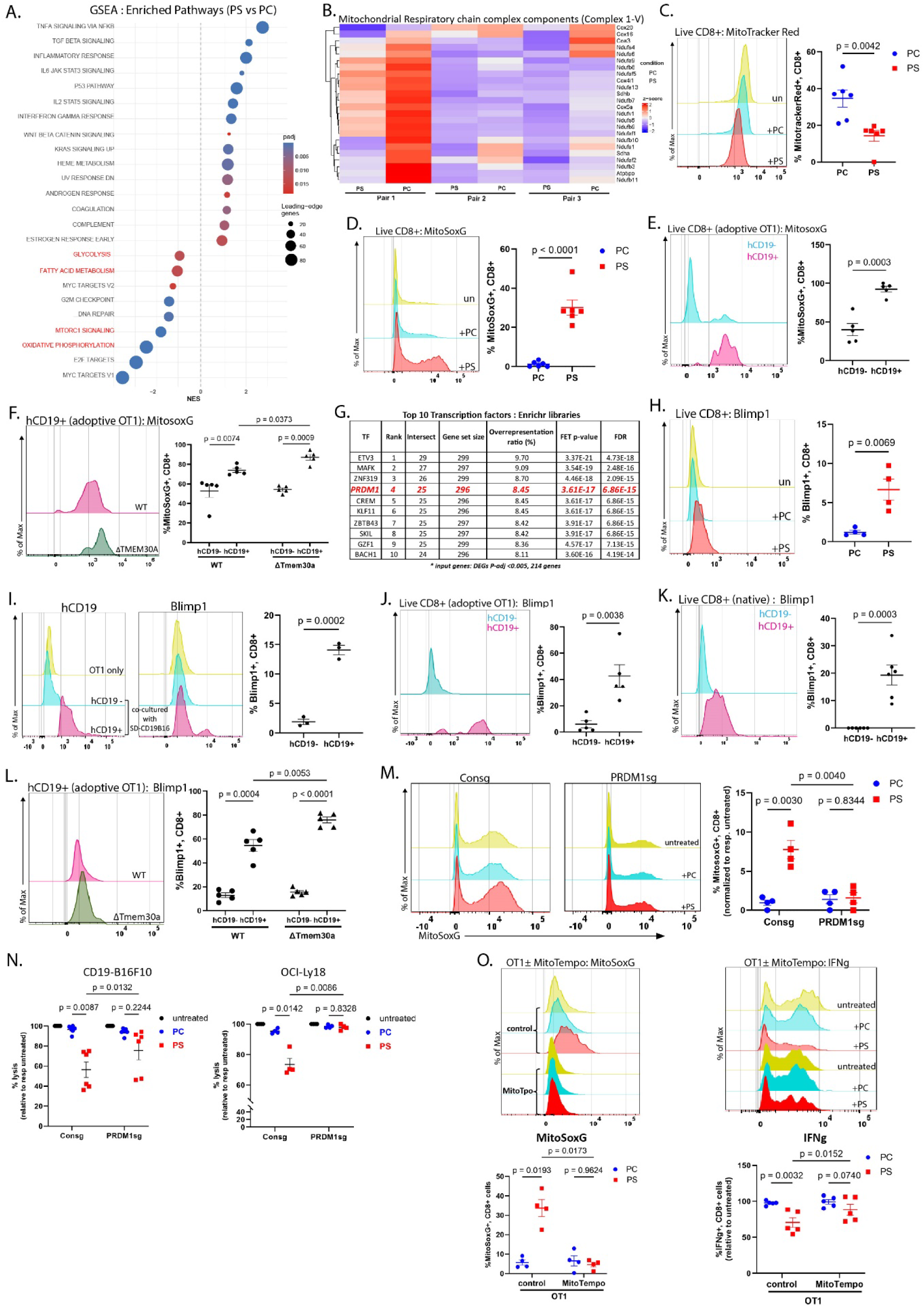
PS inactivates recipient CTLs by inducing mitochondrial dysfunction. **a)** GSEA of RNA-seq data from PS-vs. PC-liposome-treated OT-I CTLs (adjusted *P* < 0.02; *n* = 3/group, paired analysis). Metabolic pathways are shown in red; bubble size represents leading-edge gene count, and color indicates FDR *q*-value. **b)** Heatmap showing the expression of mitochondrial respiratory chain complex genes. **c)** Mitotracker Red fluorescence and %MitoRed+ CD8+ cells, and **d)** MitoSOX fluorescence and %MitoSOX+ CD8+ cells, upon PC/PS liposome treatment. **e)** MitoSOX intensity and %MitoSOX+ CD8+ cells (of live hCD19+ or hCD19-populations) in tumor-infiltrating OT-I CTLs from *Rag1*⁻/⁻ mice bearing hCD19-B16F10 (n = 5). **f)** MitoSOX intensity on live hCD19+ TI-OT-I cells, and %MitoSOX+ CD8+ (of live hCD19+ or hCD19+ populations) from wild-type or *Tmem30a*-KO hCD19-B16F10-bearing *Rag1*⁻/⁻ mice (n = 5/group). **g)** Top enriched TFs for significant DEGs (214 input genes, adjusted *P* ≤0.005), using ChEA3 (Enrichr library). **(*h-l)*** Blimp-1 expression (%Blimp-1+ CD8+, of respective live parent gates) in: **h)** PC/PS-liposome-treated OT-1 CTLs; **i)** hCD19+ vs. hCD19-OT-1 CTLs from SD-CD19B16 co-cultures (left– hCD19 transfer, right– Blimp-1); **j)** TI-OT-1 CTLs from hCD19-B16F10-tumor-bearing mice; **k)** endogenous TI-CTLs from immunocompetent hCD19-B16F10-bearing C57BL6 mice (n=6/group); and **l)** TI-OT-1 CTLs from WT or Tmem30a-KO hCD19-B16F10-bearing Rag-/-mice (histogram: Blimp-1 intensity on hCD19+ CD8+ CTLs, n=6/group). **m)** MitoSOX intensity in PC/PS-treated control or *PRDM1*-KO human CD8+ T-cells. **n)** Cytotoxic ability of PC/PS-treated control or *PRDM1*-KO human CAR-T cells towards-i) hCD19-B16F10 and ii) OCI-ly18. **o)** MitoSOX (left) and IFNg (Right) in ± MitoTEMPO OT-1 cells treated with PC/PS liposomes (%marker+ CD8+, normalized to respective untreated controls). Data represents 3-6 independent experiments (mean ± SEM); unpaired two-tailed t-test ***(c-e, h-k)*** or two-way ANOVA ***(f, l-o)***.

These results suggest that PS may undermine mitochondrial function in recipient CTLs. We next focused on this possibility because continuous antigen stimulation and hypoxia are known to drive T-cell exhaustion by altering their metabolic programming and subsequently inducing mitochondrial failure ^27^. Given the central role of mitochondrial integrity in CTL survival, and the marked increase in surface PS on intratumoral CTLs ^24,25,28^, we hypothesized that the acquisition of tumor-derived PS via blaptotrogocytosis alters the metabolic state of recipient CTLs and drives their inactivation.

Consistent with this hypothesis, PS liposome treatment of OT-1 CTLs elicited mitochondrial dysfunction characterized by decreased mitochondrial membrane potential (Fig 5C) and elevated mitochondrial reactive oxygen species (mtROS) levels **(Fig 5D**). Increased mtROS were similarly detected in PS-treated human CAR T cells (**S6A**). TIM-3 inhibition attenuated mtROS accumulation in both human CAR T cells and mouse OT-I CTLs treated with PS liposomes (**S6A-B**). Importantly, CTLs from hCD19-B16F10 tumors (**Fig 1G**) showed that specifically blaptotrogocytosis-experienced (CD19+) T cells exhibited significantly higher levels of mtROS than CD19-CTLs (**Fig. 5E**). Likewise, adoptive transferred CD19+ OT-I CTLs from the TMEMly30a-deficient tumors displayed further increased mtROS levels relative to CD19+ CTLs isolated from control tumors (**Fig. 5F**). Blaptotrogocytosis-induced metabolic dysregulation was further supported by metabolomics which revealed that TRG+ CTLs accumulate oxidized fatty acids and sphingolipids (**Fig. 2A**).

To further identify the key upstream regulators governing this PS-induced metabolic dysfunction, we performed transcription factor enrichment analysis using ChEA3 platform on our top differentially expressed genes with a stringent cutoff of adjusted p < 0.005. This unbiased computational screen identified the top 5 upstream regulators within the Enrichr co-occurrence library, including Etv3, Mafk, Znf319, Prdm1, and Crem (**Fig. 5G**). From this list, we prioritized focusing on *Prdm1* (gene encoding the Blimp1 transcription factor, rank#4; p-value = 3.61 x 10^-17^) because of its established biological roles in metabolic reprogramming ^27^ and in driving CD8+ T cell exhaustion ^29^. Consistent with our analysis predicting Blimp1 as an upstream regulator, Blimp1 protein expression was indeed significantly increased following PS liposome treatment in OT-1 CTLs (**Fig. 5H**). Blaptotrogocytosis-experienced CD19+ OT-1 cells co-cultured with serum-deprived CD19-B16 cancer cells similarly showed elevated Blimp1 expression **(Fig. 5I**). *In vivo*, adoptively transferred CD19+ OT-I CTLs (**Fig. 5J**) and endogenous CD19+ TILs (**Fig. 5K**) isolated from CD19-B16 tumors likewise showed significantly higher Blimp1 levels than their CD19-counterparts. Furthermore, CD19+ OT-I CTLs recovered from *Tmem30a*-deficient tumors expressed higher Blimp1 levels than the CD19+ OT-1 CTLs from the control tumors (**Fig. 5L**).

To test whether BLIMP1 is functionally required for PS-induced CTL dysfunction, we knocked out *PRDM1* in human anti-CD19 CAR T cells (**S6C**). Importantly*, PRDM1*-deficient CTLs retained their ability to acquire surface PS (**S6D**), and BLIMP1 induction preceded mtROS accumulation following PS exposure (**S6E**), implicating BLIMP1 as a downstream mediator of PS-induced metabolic dysfunction. Genetic ablation of *PRDM1* protected CAR T cells from PS-induced mtROS accumulation (**Fig. 5M**) and preserved their anti-cancer cytotoxic capacity *in vitro* (**Fig. 5N**), establishing that BLIMP1 is pivotal for PS-driven mitochondrial dysfunction and CTL inactivation.

To test whether scavenging mtROS could potentially rescue PS-induced exhaustion, we treated CTLs with MitoTEMPO, a mitochondrial-targeted superoxide dismutase mimetic ^30–32^. Treatment with MitoTEMPO prevented mtROS accumulation in CTLs exposed to PS-liposomes **(Fig. 5O, left)** or *Tmem30a*-deficient CD19-B16F10 cancer cells **(S6F**). Importantly, MitoTEMPO-treated CTLs retained IFN-γ expression even in response to PS exposure **(Fig. 5O, right**). Collectively, these findings show that blaptotrogocytosis-mediated PS transfer inactivates CTLs through BLIMP1-dependent mitochondrial dysfunction and oxidative stress, which can be rescued by targeting mitochondrial ROS.

### The PS–TIM-3–BLIMP1 axis is associated with clinical failure of CAR T therapy and restricts CAR T-cell anti-tumor efficacy in mouse model of lymphoma

A previous study has shown that deletion of TMEM30A in cancer cells correlates with improved patient survival in diffuse large B cell lymphoma (DLBCL) ^33^. We evaluated the clinical relevance of *TMEM30A* expression in skin cutaneous melanoma (SKCM), by performing a multivariate Cox regression analysis, adjusting for tumor purity as a confounding covariate using the TIMER3 platform. Elevated *TMEM30A* expression was independently associated with a significant survival advantage, implicating a protective role for TMEM30A in SKCM independent of immune cell infiltration density (**Fig 6A**). Additionally, an immune cell infiltration analysis revealed that increased TMEM30A expression in tumors correlated with increased CD8+ T cell infiltration within the SKCM microenvironment (**S6G**).

**Figure 6.**
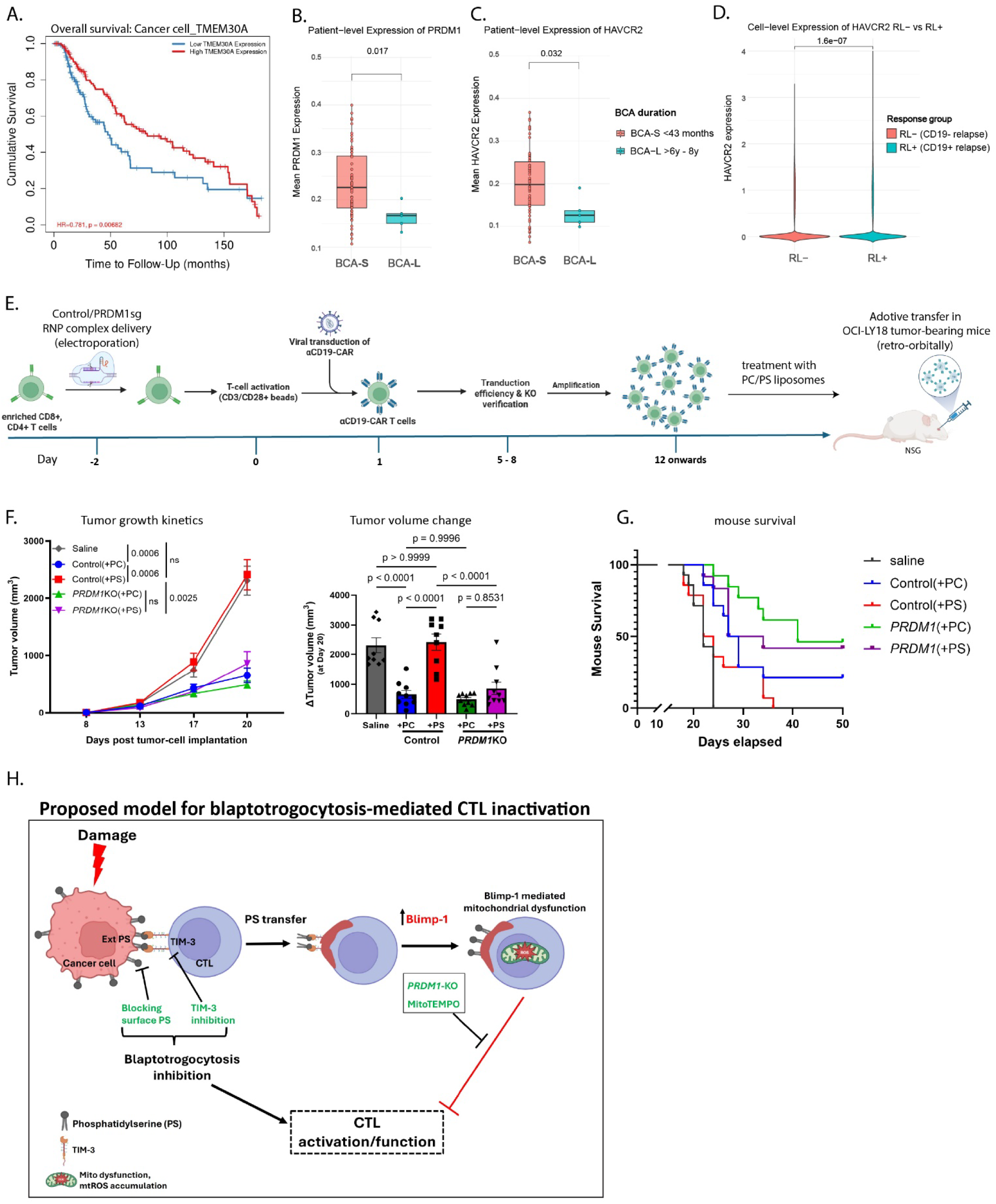
Modulation of blaptotrogocytosis regulators affects anti-tumor efficacy of CAR T cells. **a)** Kaplan–Meier analysis of *TMEM30A* expression and overall survival in patients with SKCM (TIMER3; n = 444; log-rank P = 0.0068; HR = 0.781). ***(b-c)*** Normalized basal expression of **b)** *PRDM1* and **c)** *HAVCR2* in pre-infusion αCD19 CAR T cells from pediatric ALL patients with short-term (BCA-S, n = 66) or long-term (BCA-L, n = 5) BCA; dots indicate individual patients (two-tailed Wilcoxon rank-sum test, unadjusted P = 0.017 and 0.032; BH-adjusted P = 0.0167 and 0.0319). **d)** Single-cell *HAVCR2* expression in CAR-T cells from patients with CD19-negative (RL−, n = 117,491 cells) or CD19-positive (RL+, n = 97,300 cells) relapsed disease (two-tailed Wilcoxon rank-sum test, P = 1.60 × 10⁻⁷). **e)** Experimental schema testing PS effect and *PRDM1* deficiency on CAR-T efficacy *in vivo*. **f)** Tumor growth kinetics (left; two-way ANOVA with multiple comparisons) and ΔTumor volume at day 20 post-treatment (right; one-way ANOVA), mean± SEM, n ≥ 9-10 mice/group. **g)** Kaplan-Meier survival curves (2 experiments, n ≥ 14 mice/group, Mantel-Cox log-rank p-value <0.0001; median survival (days): saline – 22, NTC(+PC) – 28, NTC(+PS) – 23, BKO(+PC) – 41 and BKO(+PS) – 31. **h)** Proposed model for blaptotrogocytosis-mediated CTL inactivation.

Next, to assess the translational relevance of TIM-3-dependent blaptotrogocytosis and PS-driven BLIMP1 induction, we analyzed gene expression data from the pre-infusion anti-CD19 CAR T-cell products from pediatric patients with acute lymphoblastic leukemia (ALL) to evaluate the correlation between *HAVCR2* (TIM-3) and *PRDM1* (BLIMP1) expression with long-term clinical persistence ^34^. In this dataset, CAR T cell functional persistence - manifested clinically by the duration of B cell aplasia (BCA) - inversely correlated with *PRDM1* expression.

Specifically, pre-infusion manufactured CAR T-cells from patients with short-term B cell aplasia (BCA-S, defined as <43 months, combining BCA-1, −2, and −3 cohorts), including non-responders and patients with relapsed disease, showed significantly higher basal *PRDM1* expression (**Fig 6B**) compared to long-term B cell aplasia (BCA-L, cohorts with 82-106 months of BCA duration). This finding links Blimp1 with CAR T-cell dysfunction. Consistent with our model, CAR T-cell products from the BCA-S cohort displayed a concurrent increase in TIM-3 (HAVCR2) expression **(Fig 6C**) relative to BCA-L products. Notably, single-cell analysis further revealed that *HAVCR2* transcripts were significantly enriched within CAR T cells mapping to the CD19-positive relapse (RL+) cohort compared with those from patients with CD19-negative relapse (RL-) (**Fig 6D**).

We next sought to model these observations *in vivo,* using an adoptive transfer model wherein immunocompromised NSG mice bearing human DLBCL OCI-Ly18 solid tumors were treated with human anti-CD19 CAR T cells (**Fig 6E**). As expected, adoptive transfer of non-targeting control (NTC) CAR T cells pretreated with control PC liposomes efficiently controlled tumor growth (**Fig 6F**) and significantly extended animal survival (**Fig 6G**). In contrast, pretreatment of NTC CAR T cells with PS liposomes drastically attenuated their anti-tumor efficacy, resulting in accelerated tumor growth (**Figure 6F-G**).

Previous studies have demonstrated that knockout of *PRDM1* increases the anti-tumor efficacy of CAR T cell adoptive transfer in NALM-6 leukemia model ^35^. In our DLBCL OCI-Ly18 model, *PRDM1* knockout (BKO) provided baseline therapeutic benefit without significantly altering clearance relative to the NTC control. However, this knockout rendered CAR T cells resistant to PS-induced suppression (**Fig. 6F-G**). Overall, these *in vivo* studies demonstrate that *PRDM1* deficiency rescues CAR T cells from PS-induced metabolic exhaustion and functional inactivation, confirming a crucial role for the PS-BLIMP1 axis in mediating blaptotrogocytosis-driven immune evasion.

## Discussion

Here we describe blaptotrogocytosis as an antigen-independent PS-TIM-3-driven transfer of plasma membrane patches from damaged cancer cells onto adjacent effector CTLs, leading to BLIMP1-mediated mitochondrial dysfunction and inactivation of these CTLs (**Fig 6H**). In addition to known roles of externalized PS in promoting plasma membrane repair at the cellular level ^36,37^ and in efferocytosis-associated immunosuppressive effects^12^, PS-driven blaptotrogocytosis likely adds an important layer of mechanisms protecting the damaged tissue from CTL-elicited immunopathology and may function as a major regulator of non-specific peripheral tolerance. Additional studies are needed to determine whether blaptotrogocytosis also involves and affects recipient natural killer cells.

Blaptotrogocytosis and resulting acquisition of PS by CTLs appears to provide a shortcut mechanism for achieving CTL exhaustion, which otherwise relies on upregulation of PS synthesis genes in CTLs, as was demonstrated under conditions of chronic viral infection^24^. In the latter model, increased PS biosynthesis and resulting PS overload in CTLs also led to inactivation of the antigen-presenting cells^24^. Future studies should examine the importance of such additional indirect mechanisms in the immunosuppressive effects of blaptotrogocytosis.

It must be noted that certain mechanistic complexities likely contribute to non-specific inactivation of CTLs adjacent to a cancer cell in the context of blaptotrogocytosis. First, in the absence of cell injury, PS can become externalized in response to cell invasion and stress pathways activated in the TME ^11,38^. Second, PS can engage other receptors besides TIM-3 ^12–15^. Finally, TIM-3 can get activated by other ligands besides PS and can inhibit T cell function in other manners besides acquisition of PS that may involve Fyn kinase-dependent inhibitory phosphorylation ^39^. Such mechanistic redundancies might contribute to somewhat underwhelming clinical responses to inhibitors of PS or TIM-3^40–42^.

Although PS itself can efficiently suppress CTL functions, we should not rule out the importance of PS-TIM-3-dependent transfer of additional biomolecules, including other types of lipids, that can contribute to CTL exhaustion and inactivation. While CTLs require lipids for their normal function, an excess of lipids in these cells can disrupt the homeostasis of the endoplasmic reticulum (ER), triggering lipogenic ER stress responses ^43–45^. For example, a surplus of long-chain fatty acids in CD8⁺ T cells elicits mitochondrial dysfunction and impairs their effector function^20^. An overload of cholesterol promotes the ER stress-driven upregulation of PD-1 and ensuing exhaustion of CTLs ^19^. Transfer of cholesterol from the cancer-associated fibroblasts onto CTLs also inhibits their function in the context of the pancreatic adenocarcinoma TME ^11^. Finally, oxidized lipids transferred onto CTLs can also suppress their activation and function ^22,23^. In addition, we cannot rule out immunosuppressive mechanisms elicited by some of the plasma membrane proteins transferred onto CTLs during blaptotrogocytosis.

While protecting injured tissues from CTL-driven immunopathology, blaptotrogocytosis may represent a grave problem for anti-cancer therapies. The demonstrated ability of damaged cancer cells to suppress intratumoral CTLs prompts an uncomfortable realization that even a successful infliction of injury intrinsically undermines tumor clearance by CTLs and may lead to acquired resistance. Future studies are required to assess a potential contribution of this phenomenon to the lack of added benefit for patients receiving immunotherapy by adding the chemotherapeutic regimens ^46^.

Two approaches may address this problem. First, temporal segregation or staggering of cytotoxic events driven by chemotherapeutics with CTL reactivation via immune checkpoint blockade or CAR T cell transfer may prove advantageous for patients receiving combination treatments. Second, development of potent blaptotrogocytosis inhibitors that are clinically superior to currently available inhibitors of PS or TIM-3 is warranted.

## Required statements

### Author Contributions

**MRD:** Study design, Investigation, Data curation, Methodology, Formal analysis, Validation, Visualization, Writing-original draft preparation; **QTB, AH, ZL, MY, EB:** Methodology, Data curation; **AS, AlW, LJ:** Data curation, Bioinformatic analysis; **AaW, KT, MJA, IAA, YF, CK, JAD, WG, VSS:** Conceptualization, Supervision, Writing-review C revision, Resources; **SYF:** Conceptualization, Supervision, Study design, Funding acquisition, Writing-review C revision, Project administration. All authors have reviewed and approved the final version of this manuscript.

### Data Availability

The RNA-seq data generated in this study have been deposited in the Gene Expression Omnibus (GEO) under accession code GSE355206. The processed lipidomics data and all other data supporting the findings of this study are available within the paper and its Supplementary Information files. Raw mass spectrometry data files are available from the corresponding author upon reasonable request. Source data are provided with this paper.

## Acknowledgements

We thank Dr. Ellen Puré and the members of Guo, Fan, and Koumenis labs for discussion and criticism. Penn Cytomics and Cell Sorting Shared Resource Laboratory at the University of Pennsylvania (RRID:SCR_022376). Penn Cytomics is partially supported by the Abramson Cancer Center NCI Grant (P30 016520). We are also thankful to Gordon Ruthel and The Cell C Developmental Biology (CDB) Microscopy Core (RRID SCR_022373) for help with confocal microscopy; and Tapan Ganguly, Erik Toorens, John Tobias and the Penn Genomics and Sequencing Core for help with RNA-sequencing (RRID:SCR_024999).

## Methods and Materials

### Experimental models

#### Cancer cell lines

Murine melanoma B16F10 cells were obtained from ATCC (CRL-6475), and murine colon adenocarcinoma MC38 cells were purchased from Kerafast (ENH204-FP). OVA-expressing MC38 (OVA-MC38) cells were a gift from S. Ostrand-Rosenberg (University of Maryland), and human CD19-expressing B16F10 (hCD19-B16F10) cells were kindly provided by A. Minn (University of Pennsylvania) ^47^. Stable luciferase-expressing reporter lines were generated by transducing OVA-B16F10 and hCD19-B16F10 cells with MSCV-Hygro-Luc retrovirus, followed by hygromycin selection; positive clones were validated using the ONE-Glo Luciferase Assay System (Promega, E6110) according to the manufacturer’s instructions. Luciferase-expressing human diffuse large B-cell lymphoma (DLBCL) OCI-Ly18 cells were kindly provided by M. Ruella (University of Pennsylvania) ^48^. All B16F10- and MC38-derived lines were maintained in Dulbecco’s modified Eagle’s medium (DMEM; Gibco, 11965092) supplemented with 10% (v/v) heat-inactivated fetal bovine serum (FBS; Gibco) and 100 U ml⁻¹ penicillin–streptomycin (Corning). OCI-Ly18 cells were cultured in RPMI 1640 (Corning, 10-041-CV) supplemented with 10% (v/v) FBS and 100 U ml⁻¹ penicillin–streptomycin.

#### Generation of *Tmem30a*-knockout cell lines

*Tmem30a*-knockout (*Tmem30a*-KO) MC38, B16F10, and hCD19-B16F10 cell lines were generated by transient transfection with all-in-one CRISPR–Cas9 plasmids targeting murine *Tmem30a* (GeneCopoeia, MCP304097-CG12-3-B) or a scrambled sgRNA control (GeneCopoeia, CCPCTR01-CG12-B) using Lipofectamine 2000 (Invitrogen) according to the manufacturer’s instructions. Briefly, cells were seeded in 6-well plates to reach ∼70% confluency the next day. Before transfection, culture medium was replaced with 500 µl serum- and antibiotic-free Opti-MEM (Life Technologies, 31985062). For each well, 4 µg plasmid DNA diluted in 250 µl Opti-MEM was incubated for 5 min at room temperature and combined with 20 µl Lipofectamine 2000 diluted in 250 µl Opti-MEM, followed by gentle mixing and a 20 min incubation at room temperature. The 500 µl DNA–lipid complex was added dropwise to cells; 6 h later, 1 ml complete DMEM was added, and cells were switched back to fresh complete DMEM (10% [v/v] FBS, 100 U ml⁻¹ penicillin–streptomycin) the following day. At 48 h post-transfection, cells were selected with G418 (neomycin; 2 mg ml⁻¹) for 5 days. We confirmed Tmem30a deletion at the protein level by immunoblotting and functionally validated it by assessing constitutive surface phosphatidylserine exposure via Annexin V staining (Supplementary Fig. 2e).

All parental and engineered cell lines routinely tested negative for *Mycoplasma* contamination (MycoAlert Mycoplasma Detection Kit, Lonza) and were used within 1 month of thawing for experiments.

#### Mouse models

All animal housing and experimental procedures were approved by and conducted in accordance with the Institutional Animal Care and Use Committee (IACUC) at the University of Pennsylvania (protocol no. 803995). Six-to eight-week-old C57BL/6J wild-type mice (The Jackson Laboratory, 000664; RRID:IMSR_JAX:000664) and 6–7-week-old *Rag1*⁻/⁻ mice (C57BL/6J-*Rag1*ᵉᵐ¹⁰ᴸᵘᵗᶻʸ/J; The Jackson Laboratory, 034159) were purchased from The Jackson Laboratory. Seven-to eight-week-old and 15–16-week-old NOD.Cg-*Prkdc*ˢᶜⁱᵈ *Il2rg*ᵗᵐ¹ᵂʲˡ/SzJ (NSG) mice (The Jackson Laboratory, 005557) were used for the young and older adult *PRDM1*-KO CAR-T therapy experiments, respectively. OT-I TCR-transgenic mice (C57BL/6-Tg(*TcraTcrb*)1100Mjb/J; The Jackson Laboratory, 003831) ^49^ were originally obtained from The Jackson Laboratory and bred in-house; homozygous mice (8-12 weeks old) were used to maintain the breeding colony; and 6-12 weeks old mice were used as T-cell donors. Both male and female mice were used to generate OT-1 cells.

## Methods

### OT1 CD8+ T-cell generation

OT-I CD8⁺ T cells were isolated from the spleens of 7–12-week-old OT-I mice. Briefly, excised spleens were washed once with T-cell medium consisting of RPMI 1640 (Corning, 10-041-CV) supplemented with 10% (v/v) heat-inactivated FBS and 100 U ml⁻¹ penicillin– streptomycin and mechanically dissociated through a 70 µm cell strainer using the plunger of a 5 ml syringe to obtain a single-cell suspension. Cells were pelleted by centrifugation at 500*g* for 5 min and resuspended in 1× RBC Lysis Buffer (eBioscience, 00-4300-54) for 5 min at room temperature. After quenching and washing with complete T-cell medium, cells were centrifuged at 500*g* for 5 min and cultured in complete T-cell medium supplemented with 55 µM β-mercaptoethanol (1:1,000 dilution; Gibco) and 1 µg/ml ovalbumin peptide (OVA₂₅₇–₂₆₄; Sigma-Aldrich, S7951) for 48 h. Activated OT-I CD8⁺ T cells were used for all subsequent in vitro blaptotrogocytosis, antigen-driven trogocytosis, and liposome assays, as well as adoptive transfer into *Rag1*⁻/⁻ mice.

### Trogocytosis C blaptotrogocytosis assays

**To assess antigen-driven trogocytosis**, parental, OVA-expressing B16F10 or MC38, or hCD19-expressing B16F10 cancer cells were membrane-labeled using the PKH67 Green Fluorescent Cell Linker Kit (Sigma-Aldrich, MINI67). Briefly, adherent cancer cells were detached using non-enzymatic Cellstripper (Corning, 25-056-CI) and washed twice with PBS to remove residual serum. Cell pellets were resuspended in 500 µl PKH67 working dye solution (1:750 dilution in Diluent C) and incubated for 10 min at room temperature in the dark. Staining was quenched by adding complete DMEM (10% [v/v] FBS, 100 U ml⁻¹ penicillin–streptomycin) for 2–3 min. Cells were pelleted by centrifugation (500*g*, 5 min), washed twice with complete DMEM, and resuspended in complete medium for co-culture. PKH67-labeled OVA-B16F10 or OVA-MC38 cells were co-cultured with OT-I CTLs, and hCD19-B16F10 cells were co-cultured with mouse or human αCD19 CAR-T cells, at an effector-to-target (E:T) ratio of 5:1 (1:5 cancer cell-to-T-cell ratio) in round-bottom 96-well plates for 15 min at 37 °C in 5% CO₂. Parental B16F10 or MC38 cells co-cultured with OT-I CTLs or αCD19 CAR-T cells served as antigen-negative controls. After 15 min, plates were centrifuged (500*g*, 5 min, 4 °C) and cells were immediately processed for flow cytometry staining.

**To examine damaged-induced, blaptotrogocytosis**, cancer cells were co-cultured with antigen-non-specific CTLs. Parental B16F10 or MC38 cells were membrane-labeled with PKH67 as described above to assess membrane transfer, whereas hCD19-expressing B16F10 (hCD19-B16F10) cells were used to assess protein transfer. PKH67-labeled or hCD19-B16F10 cancer cells were subjected to serum deprivation (18 h) or treated with the ferroptosis inducer erastin (20 µM, 1 h) or the chemotherapeutics etoposide (250 µM, 1 h) or cisplatin (500 µM, 1 h), followed by washing with complete medium before co-culture. For UV-induced damage, PKH67-labeled cells were exposed to 40 mJ cm⁻² UV irradiation using a GS Gene Linker UV Chamber (Bio-Rad) according to the manufacturer’s instructions and rested for 1 h before co-culture. Damaged PKH67-labeled parental B16F10 or MC38 cells were co-cultured with OT-I, mouse αCD19 CAR-T, or human αCD19 CAR-T CTLs, whereas damaged hCD19-B16F10 cells were co-cultured with OT-I CTLs at a 1:5 cancer cell-to-T-cell ratio in round-bottom 96-well plates for 15 min at 37 °C in 5% CO₂. Cells were then pelleted (500*g*, 5 min, 4 °C), washed twice with PBS, and stained for flow cytometry to quantify PKH67+ membrane or hCD19 protein transfer as well as surface (PD-1, CTLA-4, LAG-3) and intracellular (TOX, Blimp-1) T-cell exhaustion markers. In an orthogonal model, human OCI-Ly18 cells endogenously expressing CD19 were serum-starved for 18 h and co-cultured with OT-I CTLs (1:5 cancer cell-to-CTL ratio) for 15 min to measure human CD19 transfer onto OT-I CTLs.

### Flow staining

Post co-culturing, centrifuged cells were washed twice with PBS, and blocked for non-specific binding using either anti-mouse CD16/CD32 (BD Biosciences, AB_394656) when staining OT-1/mCARTs or anti-human BD Fc block while staining hCARTs (BD Biosciences, cat#564220), and stained with live/dead fixable viability dye Alexa Fluor 780 (Invitrogen, cat#65-0865-14) (15 mins at RT in dark). Following Fc block and live/dead staining, cells were washed once with FACS buffer (PBS with 2% FBS) and stained with the indicated antibodies at room temperature, for 15 mins in dark (at 1:100 dilution). Stained cells were washed thrice, resuspended in FACS buffer, and immediately analyzed by flow cytometry for membrane transfer and other cell surface markers. All the washing steps post-coculture, expect for live/dead and antibody staining were performed on ice, cells were kept on ice till running on the flow cytometer. For intracellular markers (TOX, Blimp1), following cell-surface staining, cells were fixed, permeabilized, and stained using the FoxP3/Transcription factor staining buffer kit, (Invitrogen, cat.# 00-5523) as per manufacturer’s instructions. To assess effector molecules or exhaustion marker expression specifically on trogocytosis-experienced (TRG+) versus trogocytosis-naive (TRG-) CTLs within the same co-cultures, live CD8+ T cells were gated into hCD19+ vs. hCD19- or PKH67+ vs. PKH67-subpopulations for downstream marker analysis.

Surface-exposed phosphatidylserine (PS) was detected using an Annexin V staining protocol (BioLegend) according to the manufacturer’s instructions. Briefly, following viability dye and surface antibody staining, cells were washed twice and resuspended in 1X Annexin V Binding Buffer (BioLegend, 422201) at 1 × 10⁶ cells/ml. Cells were incubated with fluorochrome-conjugated Annexin V (5 µl per 100 µl cell suspension; BioLegend) for 15 min at room temperature in the dark. After staining, cells were washed once, diluted with Annexin V Binding Buffer, and immediately acquired by flow cytometry.

### Characterization of blaptotrogocytosis

For time-course kinetics, control, cisplatin-treated, or serum-deprived (SD) PKH67-labeled B16F10 cells were co-cultured with OT-I CTLs for 15, 30, 60, or 120 min, and PKH67⁺ membrane transfer was quantified by flow cytometry as described above. To determine whether membrane transfer is an active, temperature-sensitive process, control, UV-irradiated, or erastin-treated PKH67-labeled B16F10 cells were co-cultured with OT-I CTLs at either 4 °C or 37 °C for 15 min, followed by flow cytometric quantification of PKH67⁺ transfer. To evaluate contact dependence, control or Cisplatin-treated or SD CD19-B16F10 cells were cultured with OT-I CTLs either in direct contact within a single well or physically separated in 24-well Transwell plates (0.4 µm pore size; Corning Costar #3470), with hCD19-B16F10 cells seeded in the lower chamber and OT-I CTLs in the upper insert, for 15 min to 4 h; OT-I CTLs were then harvested from the upper inserts or direct co-cultures and analyzed by flow cytometry for hCD19 protein transfer. To assess the requirement for actin polymerization and cytoskeletal reorganization, OT-I CTLs were pretreated with vehicle control or latrunculin B (2µM, 30 min; Sigma-Aldrich, 428020) prior to co-culture with control or damaged PKH67-labeled B16F10 cells. To test the requirement for membrane fusion, OT-I CTLs were pretreated with vehicle control or 25-hydroxycholesterol (25HC; 6µM, 2h [sigma-Aldrich, H1015]) prior to co-culture with control or damaged PKH67-labeled B16F10 cells.

### Liposome treatment

To assess direct PS acquisition, CTLs (OT-1, mCART19 or hCART19) were incubated with 50µM PC or PS liposomes (Sigma-Aldrich, CPS-513 or CPS-514) for 20 minutes at 37^0^ C, washed twice with PBS, and stained with Annexin V. Surface PS levels were quantified on live CD8+ T cells.

To evaluate the effect of PS on T-cell effector function, CTLs (OT-1, mCART19 or hCART19 or human CD8+ T-cells) were treated with 50µM control PC or PS liposomes for 18 hours, with a protein transport/Golgi inhibitor (BD Biosciences, 554724) during the last 3 hours. Cells were then stimulated with cognate antigen-expressing target cells (OVA-B16F10 for OT-I CTLs or hCD19-B16F10 for αCD19 CAR-T cells or PMA/ionomycin for non-transduced human CD8⁺ T cells) for 30 min prior to intracellular cytokine staining.

To disrupt PS–TIM-3 interactions, donor cancer cells (antigen-expressing or damaged) were pretreated with anti-PS antibody (10 µg/ml; Sigma-Aldrich [05-719, clone 1H6]), recombinant Annexin V (20 µg per 10⁶ cells ml⁻¹ [BD Pharmingen, cat#556416]), or the scramblase inhibitor niclosamide (2µM, 3h [vendor, cat#50-65-7]) prior to co-culture. To pharmacologically inhibit TIM-3 on recipient OT-I CTLs, splenocytes were cultured in medium containing the TIM-3 inhibitor ML-T7 (5 µM [MedChemExpress, cat#HY-163028]) starting at day 0 alongside OVA₂₅₇–₂₆₄ peptide stimulation for 48 h prior to assays. Mouse or human αCD19 CAR-T cells were pretreated with ML-T7 (5 µM) for 48 h prior to trogocytosis, blaptotrogocytosis, or liposome-treatment experiments. Antibody-mediated TIM-3 blockade was achieved by pretreating OT-I or mouse CAR-T cells with anti-TIM-3 blocking antibody (15 µg ml⁻¹ [BioXCell, Cat#BE0115, clone: RMT3-23]) for 30 min prior to co-culture and maintaining the antibody throughout the co-culture period.

### Cytotoxicity assays

OT-I CTLs pretreated with PC or PS liposomes (50 µM, 18 h) were co-cultured with adherent firefly luciferase-expressing OVA-MC38 or OVA-B16F10 target cells [at an E:T ratio of 1:5] for 8 h and 24 h, respectively. Following co-culture, target cell viability was determined by measuring bioluminescence using the ONE-Glo Luciferase Assay System (Promega, E6110) according to the manufacturer’s instructions. Specific target cell lysis (%lysis) was calculated from residual bioluminescence intensity relative to target cells co-cultured with untreated OT-I CTLs. Similarly, the cytolytic activity of controlsg (control) or *PRDM1*-KO (BKO) human αCD19 CAR-T cells pretreated with PC or PS liposomes (50 µM, 18 h) was assessed by co-culture with luciferase-expressing hCD19-B16F10 or OCI-Ly18 target cells [at an E:T ratio of 5:1] for 12 h and 6 h, respectively, followed by bioluminescence measurement as described above. Formulae for calculating %lysis = target T cell (no T-cell) - Test sample (i.e., +)/ target cell (no T-cell) - maximum killing (i.e., with H2O) * 100

### Assessment of mitochondrial membrane potential and reactive oxygen species

Mitochondrial membrane potential was assessed using MitoTracker Red FM (Invitrogen, M46751). CTLs treated with PC or PS liposomes (50 µM, 1.5 h) or co-cultured with cancer cells (1 h) were washed and stained for viability and surface CD8 (without fixation). Cells were then subsequently incubated with MitoTracker Red FM (200 nM in pre-warmed HBSS) for 30 min at 37 °C in 5% CO₂ protected from light, according to the manufacturer’s instructions, and immediately analyzed by flow cytometry, gating on live CD8⁺ T cells.

Mitochondrial superoxide accumulation (mtROS) was measured using MitoSox Green Mitochondrial Superoxide Indicator (Invitrogen, M36005). Following PC/PS liposome treatment (18 h), or tumor dissociation, single-cell suspensions were washed and stained for live/dead dye, followed by surface staining for CD8a. Cells were then incubated with MitoSoxG working solution (5 µM in warm HBSS containing Ca²⁺ and Mg²⁺) for 30 min at 37°C in 5% CO₂ protected from light. Cells were washed twice with warm HBSS, and immediately acquired by flow cytometry, quantifying MitoSoxG Green fluorescence intensity and % MitoSoxG+ cells within the live CD8+ (and hCD19+ or hCD19-, where indicated) gates.

For mitochondrial ROS scavenging, OT-I CTLs were pretreated with MitoTEMPO (10µM for 1h, Sigma-Aldrich, Cat# SML0737) and subsequently incubated with PC or PS liposomes (18h) or co-cultured with wild-type (WT) or *Tmem30a*-KO CD19-B16F10 cells (4h), followed by flow cytometric analysis of MitoSOX Green accumulation and intracellular IFNγ expression.

### In vivo trogocytosis and blaptotrogocytosis models

To evaluate antigen-independent blaptotrogocytosis in vivo, 6–7-week-old male *Rag1*⁻/⁻mice were subcutaneously inoculated in the right flank with 5 × 10⁵ hCD19-B16F10 cells [in 100 µl sterile PBS]. Once tumors reached ∼9–10 mm^3^ in volume, mice received an intravenous adoptive transfer (via the retro-orbital sinus) of 48 h peptide-activated OT-I CD8⁺ T cells (2 × 10⁶ cells per mouse). At 18 h post-transfer, tumors and spleens were harvested and processed for flow cytometry. To examine blaptotrogocytosis and its functional consequences on endogenous CD8⁺ CTLs, 6–8-week-old male C57BL/6J mice were subcutaneously inoculated with 5 × 10⁵ hCD19-B16F10 cells, and tumors and spleens were harvested for flow cytometric analysis once tumors reached ∼9–10 mm in diameter. Similarly, to evaluate constitutive phosphatidylserine externalization in vivo, 6–7-week-old male *Rag1*⁻/⁻ mice were subcutaneously inoculated with 5 × 10⁵ wild-type (WT) or *Tmem30a*-knockout (*Tmem30a*-KO) hCD19-B16F10 cells; once tumors reached ∼9–10 mm in diameter, mice were adoptively transferred via the retro-orbital route with 48 h-activated OT-I CTLs (2 × 10⁶ cells per mouse), and tumors and spleens were collected 18 h later for flow cytometry.

For tissue dissociation, excised tumors were weighed, and approximately equal masses (g) of each tumor sample were taken for downstream processing. Tumor tissues were finely minced and enzymatically digested using a collagenase‒DNAse mixture (1 mg/ml collagenase + 50 U/ml DNAse I in RPMI supplemented with 2% FBS) at 37^0^ for 30 min with gentle agitation. Digested tumor tissues and excised spleens were mechanically dissociated through 70 µm cell strainers using the plunger of a 5 ml syringe to obtain single-cell suspensions. Cells were pelleted by centrifugation (500*g*, 5 min) and resuspended in 1× RBC Lysis Buffer (eBioscience, 00-4300-54) for 5 min at room temperature for red blood cell lysis. After neutralizing the RBC lysis buffer and washing with RPMI 1640 containing 2% (v/v) FBS, 2-2.5 x 10^6^ cells per sample were stained for surface and intracellular cell markers as described above.

### CRISPR–Cas9-mediated *PRDM1* editing in human T cells

Primary human CD4⁺ and CD8⁺ T cells were isolated from peripheral blood mononuclear cells (PBMCs) of de-identified healthy donors by negative selection (RosetteSep Human T Cell Enrichment Cocktail, STEMCELL Technologies) through the Human Immunology Core at the University of Pennsylvania (Philadelphia, PA, USA), with institutional review board (IRB) approval and informed donor consent. Isolated T cells were cultured in complete human T-cell medium consisting of RPMI 1640 (Corning, 10-041-CV) supplemented with 10% (v/v) heat-inactivated FBS, 100 U ml⁻¹ penicillin–streptomycin, 55 µM β-mercaptoethanol (1:1,000 dilution; Gibco), recombinant human IL-7 ([Gibco, 20007; 10 ng/ml]), and recombinant human IL-15 (Gibco, 20015; 10 ng/ml]; Gibco). The following day, 1 × 10⁷ T cells were washed twice with Opti-MEM (Life Technologies, 31985062) and electroporated with ribonucleoprotein (RNP) complexes comprising 10 µg Alt-R *S.p.* Cas9 Nuclease V3 (IDT), Alt-R CRISPR–Cas9 sgRNA targeting *PRDM1*: 5’-CATCAGCACCAGAATCCCAG-3 or a non-targeting *AAVS1* control: 5’-CCATCGTAAGCAAACCTTAG-3’ ^50^ (IDT), and Alt-R Cas9 Electroporation Enhancer (IDT) using the P3 Primary Cell 4D-Nucleofector X Kit (Lonza, V4XP-3024; pulse code [e.g., EO-115]). Electroporated cells were immediately transferred into 1 ml pre-warmed complete medium for recovery for 48 h and subsequently activated with Dynabeads Human T-Activator CD3/CD28 (Invitrogen, 11132D) at a 1:3 cell-to-bead ratio for 24 h [followed by lentiviral transduction with the anti-CD19 CAR construct—add brief CAR transduction step if not covered in the preceding paragraph]. *PRDM1* (Blimp-1) knockout efficiency was confirmed at the protein level by immunoblotting (Supplementary Fig. 6c).

#### In vivo *PRDM1*-knockout CAR-T cell efficacy

Six-to seven-week-old male NSG mice were subcutaneously inoculated in the right flank with 3.5 × 10⁶ luciferase-expressing OCI-Ly18 DLBCL cells suspended in a 1:1 (v/v) mixture of PBS and Matrigel (Corning). Once palpable tumors formed (∼2.5–3 mm in diameter), mice were randomized into treatment groups and intravenously injected (via the retro-orbital sinus) with either sterile saline or 2 × 10⁶ human αCD19 CAR-T cells per mouse. Prior to adoptive transfer, non-targeting control (NTC) or *PRDM1*-knockout (BKO) CAR-T cells were pretreated in vitro for 16 h with 50 µM PC or PS liposomes [and washed prior to injection]. Adoptive transfers were repeated every 4 days until the saline control group reached the humane endpoint. Tumor dimensions were measured regularly with digital calipers, tumor volume was calculated as *(length × width²)/2*, and mice were monitored for overall survival. Change in tumor volume (Δ tumor volume, mm³) was calculated as the difference between tumor volume at day 20 and tumor volume at treatment initiation. In accordance with University of Pennsylvania IACUC guidelines, the maximal permitted tumor diameter of 20 mm (2.0 cm) was not exceeded in any experiment.

### Confocal Microscopy, live cell imaging

To quantify CTL–cancer cell contacts and intercellular distances, B16F10 cells were membrane-labeled with PKH67 (Sigma-Aldrich) as described above and seeded at 8 x 10^4^ cells per dish in 35 mm polymer coverslip-bottom imaging dishes (ibidi). Activated OT-I CTLs (pretreated with the TIM-3 inhibitor ML-T7 or anti-TIM-3 blocking antibody as described above, where indicated) were labeled with 10 µM CellTracker Blue CMAC (Thermo Fisher Scientific, C2110) for [15–30] min at 37 °C, washed twice, and resuspended in complete medium. PKH67-labeled B16F10 cells were treated with cisplatin (500 µM, 1 h) in the presence or absence of anti-phosphatidylserine blocking antibody (10 µg ml⁻¹), washed, and co-cultured with 4 x 10^5^ CMAC-labeled OT-I CTLs (1:5 cancer cell-to-T-cell ratio) for 15 min at 37 °C in 5% CO₂. Live-cell imaging was performed on a Leica STELLARIS confocal microscope (Leica Microsystems) equipped with a stage-top environmental chamber (37 °C, 5% CO₂) using a 40× objective [NA, oil/water immersion]. Five randomly selected, non-overlapping fields of view were acquired per dish. Using Fiji (ImageJ), the shortest Euclidean distance from the plasma membrane of each OT-I CTL to the plasma membrane of the nearest B16F10 cell was measured, and both the percentage of OT-I CTLs in direct membrane contact with cancer cells (distance = 0 µm) and the mean CTL-to-nearest-cancer-cell distance were calculated per field.

### Western blot

Western blot was performed as per the standard established protocol. Briefly, 60 µg of total protein was separated on 10% polyacrylamide/SDS gel, transferred on to PVDF membranes, and membranes were blocked with 5% (w/v) Bovine Serum Albumin (BSA) in Tris Buffered Saline TBS (+0.1% Tween-20). Primary antibodies include: TMEM30A (Invitrogen, Cat# PA5-99762), Blimp-1 (Invitrogen, Cat# 14-5963-82), β-Actin (Cell Signaling Technology Cat# 4970, RRID:AB_2223172), Vinculin (Cell Signaling Technology Cat#13901). Horseradish peroxidase-conjugated secondary antibody used: anti-rabbit IgG (Cell Signaling Technology Cat# 7074, RRID:AB_2099233).

### RNA-seq and bioinformatic analysis

Three biological replicates of PS- or PC-liposome-treated OT-I CTLs (50 µM, 1.5 h) were collected, and total RNA was extracted using RNeasy RNA isolation kit (Qiagen, cat. no.74134). RNA purity was assessed using a NanoDrop spectrophotometer (Thermo Fisher Scientific) to confirm an *A*₂₆₀/*A*₂₈₀ ratio > 1.9, and RNA integrity number (RIN) was measured using an RNA 6000 Nano Kit on a Bioanalyzer (Agilent Technologies). Illumina Stranded mRNA Prep libraries were prepared from 100ng of total RNA according to the manufacturer’s protocol. Final libraries were sequenced in 50bp paired end configuration on a NextSeq 2000 using the NextSeq™ 1000/2000 P2 XLEAP-SBS™ Reagent Kit (100 Cycles). Sequence reads were demultiplexed in Illumina BaseSpace using the BCLConvert app (v4.3.13). https://www.illumina.co/products/by-type/sequencing-kits/library-prep-kits/stranded-mrna-prep.html https://www.illumina.com/products/by-type/sequencing-kits/cluster-gen-sequencing-reagents/nextseq-1000-2000-reagents.html#tabgroup-2-tab2

### RNA-seq data processing and analysis

Paired-end reads (2 × 50 bp; 56–85 million read pairs per library) were trimmed and quality filtered with fastp (v0.22.0) using default settings. Transcript abundance was estimated with Salmon (v1.10.3) in selective-alignment mode with GC-bias correction, using a decoy-aware index of the GENCODE vM38 transcriptome (GRCm39). All later steps were run in R (v4.6.0) with Bioconductor (v3.23) packages. Transcript estimates were imported and summarized to the gene level with tximeta (v1.30.0), and genes were annotated from Ensembl release 115 with biomaRt (v2.68.0). Differential expression between PS- and PC-liposome-treated CTLs was tested with DESeq2 (v1.52.0) using a paired design (∼ condition + mouse), so the treatment effect was estimated within each of the three mice. Wald-test p-values were adjusted by the Benjamini–Hochberg method.

Gene set enrichment analysis was run with fgsea (v1.38.0) in pre-ranked mode with default parameters, ranking genes by the DESeq2 Wald statistic. Mouse genes were mapped to human orthologs from Ensembl Compara (release 115; one-to-one, one-to-many and apparent one-to-one) so that the human MSigDB (v2026.1) gene sets could be used. Where several mouse genes mapped to one human symbol, their statistics were averaged. The Hallmark and KEGG MEDICUS (C2:CP:KEGG_MEDICUS) collections were tested. Fig. 5A shows Hallmark pathways with adjusted p ≤ 0.02. Fig. S5A shows KEGG MEDICUS reference pathways with adjusted p ≤ 0.08. Bubble plots were drawn with ggplot2 (v4.0.3). For the heatmaps (Fig. 5B, S5B–C), counts were transformed with the DESeq2 variance-stabilizing transformation and the leading-edge genes were mapped back to mouse. Values were then scaled to row z-scores and plotted with ComplexHeatmap (v2.28.0), with samples ordered by mouse so that each PS sample sits next to its PC pair.

#### Bioinformatic analysis for TF enrichment determination

To identify the potential upstream transcriptional regulators, transcription factor (TF) enrichment analysis was performed using the ChEA3 platform. Significant DEGs (input genes = 214, p-adj ≤0.005) identified from RNA-seq of PS-versus PC-liposome-treated mouse OT-1 CTLs were submitted as the query gene list. Because ChEA3 primary networks are mapped to human TF targets, mouse gene symbols were automatically converted to their corresponding human orthologs by the web server. Enriched TFs were evaluated using the integrated ‘Enrichr Queries’ library, which leverages co-occurrence networks extracted from crowdsourced, mixed-species public user submissions. TFs were ranked based on statistical significance across the 1,404 TFs within the Enrichr library, identifying *Prdm1* (Blimp-1) as ranked 4^th^ overall (Fig5G, SF5D). Extracted data parameters included: Rank (absolute ranking of the TFs), Scaled rank (normalized position within the library calculated as integer rank divided by total library TFs), Intersect (number of overlapping genes), Gene set size (number of known downstream target genes), Overrepresentation ratio percentage (calculated as intersect/gene set size x 100), and FET p-values (negative logarithm of the p-value [-log10(p-value)] calculated via Fisher’s Exact Test).

#### Untargeted lipidomics using DESI-MS analysis

DESI–MS experiments were performed using a Xevo G2-XS QToF mass spectrometer (Waters, Milford, MA) equipped with a two-dimensional DESI stage source (Prosolia) under optimized acquisition parameters. Red Sharpie and black Staedtler markers were used to assess spatial calibration for positive and negative modes, respectively. The spray solvent consisted of 98% UPLC–MS grade methanol (EMSCO Fisher, Philadelphia, PA) and 2% Milli-Q water (Millipore Sigma, Burlington, MA) containing 50 pg/μL leucine-enkephalin (Waters) as a lock mass reference for negative ion mode analysis. For positive ion mode experiments, the same solvent composition was used with the addition of 0.1% formic acid (Fisher Scientific) to enhance ionization efficiency. Solvent delivery was maintained at a flow rate of 2 μL/min using a Harvard syringe pump. Raw file was converted to imzml and processed in MSI-Eagle (1) for background subtraction and peak picking. Downstream statistical analysis and hierarchical clustering heatmaps of differentially regulated lipid classes were performed within the MSI.EAGLE platform using an ANOVA with Benjamini-Hochberg false discovery rate correction (BH-FDR < 0.05) and Tukey post-hoc tests (p < 0.05).

#### ELISA for absolute PS detection

To evaluate the total PS levels in CTLs following interaction with control or stressed B16F10 cells, OT-I CTLs or human αCD19 CAR-T (hCAR-T) cells were layered over adherent B16F10 cells (mock control, serum-deprived [SD], or cisplatin-treated) and co-cultured for 30 min at 37 °C. Non-adherent OT-I and hCAR-T cells were carefully collected, washed twice with PBS, and lysed for total protein extraction [in RIPA / lysis buffer; normalized to equal protein input]. Intracellular PS concentrations (ng/ml) were quantified using a phosphatidylserine ELISA kit. (Cat#MBS288430) according to the manufacturer’s instructions. Similarly, absolute PS concentrations (ng/ml) in OT-I CTLs following liposome exposure were determined by incubating OT-I CTLs with control PC or PS liposomes (50µM, 30min), followed by washing, total protein extraction, and PS quantification by ELISA.

#### Patient data analysis

Processed single-cell RNA-seq data from pre-infusion αCD19 CAR-T cell products of pediatric patients with acute lymphoblastic leukemia (ALL) were downloaded from the NCBI Gene Expression Omnibus (GEO) under accession number GSE262072. Using R (v4.5.1) and Seurat (v5.2.0), the processed count matrix was split by patient sample and re-normalized using the NormalizeData function with default parameters. Patient-level mean normalized basal expression levels of *PRDM1* and *HAVCR2* (encoding TIM-3) were compared across clinical response groups stratified by B-cell aplasia (BCA) duration: short-term BCA (BCA-S; BCA-1, BCA-2, and BCA-3 combined, <43 months; *n* = 66 patients) versus long-term BCA (BCA-L; >6 to 8.5 years; *n* = 5 patients) (Fig. 6b, c: Each datapoint represents an individual patient). Statistical comparisons between BCA-S and BCA-L cohorts were performed using an unpaired two-tailed Wilcoxon rank-sum test (wilcox.test in R), with *P* values adjusted for multiple comparisons using the Benjamini–Hochberg method (*PRDM1*: unadjusted *P* = 0.017, BH-adjusted *P* = 0.0167; *HAVCR2*: unadjusted *P* = 0.032, BH-adjusted *P* = 0.0319). In addition, single-cell *HAVCR2* expression levels from the same dataset (GSE262072) were compared at the individual-cell level between pre-infusion CAR-T cells from patients who subsequently experienced CD19-negative relapse (RL⁻, *n* = 117,491 cells) versus CD19-positive relapse (RL⁺, *n* = 97,300 cells) using an unpaired two-tailed Wilcoxon rank-sum test (*P* = 1.60 × 10⁻⁷; Fig. 6d).

#### For TMEM30A in publicly available patient datasets

Kaplan–Meier overall survival analysis stratified by *TMEM30A* expression (high versus low) in The Cancer Genome Atlas (TCGA) skin cutaneous melanoma (SKCM) cohort (*n* = 444 patients) was performed using the TIMER3 web server (unadjusted log-rank *P* = 0.0068; unadjusted hazard ratio [HR] = 0.781). Multivariate Cox proportional hazards regression adjusting for tumor purity (Surv(OS, EVENT) ∼ TMEM30A + Purity) confirmed that elevated *TMEM30A* expression was independently associated with improved overall survival (adjusted HR = 0.795; 95% confidence interval [CI], 0.694–0.912; Cox regression *P* = 0.001; likelihood ratio test *P* < 0.001; Wald test *P* < 0.001; overall *R*² = 0.036). In addition, the association between *TMEM30A* expression and intratumoral CD8+ T-cell infiltration in the TCGA-SKCM cohort (*n* = 470 patients) was evaluated via TIMER3 using a tumor purity-adjusted partial Spearman’s rank correlation (ρ = 0.222, *P* = 0.025).

#### Statistical analysis

Statistical analyses were performed using GraphPad Prism (version 10.4) or R (v4.6.0 / v4.5.1). Unpaired two-tailed Student’s *t*-tests were used for comparisons between two groups, and one-way or two-way analysis of variance (ANOVA) with multiple-comparisons post-hoc tests was used for comparisons across three or more groups, including longitudinal tumor growth kinetics. Survival curves were generated using the Kaplan–Meier method and compared using the log-rank (Mantel–Cox) test. Non-parametric patient cohort comparisons were performed using unpaired two-tailed Wilcoxon rank-sum tests with Benjamini–Hochberg false discovery rate (FDR) correction. Data are presented as mean ± SEM and *P* < 0.05 (α = 0.05) was considered statistically significant, with exact *P* values or significance thresholds reported in the figures. No statistical methods were used to predetermine sample sizes, but sample sizes were chosen based on prior experience and standard practice in the field. Mice were randomized into treatment groups prior to therapy initiation; investigators were not blinded to group allocation during experiments and outcome assessment.

## Supplementary figures

**Supplementary Figure 1.**
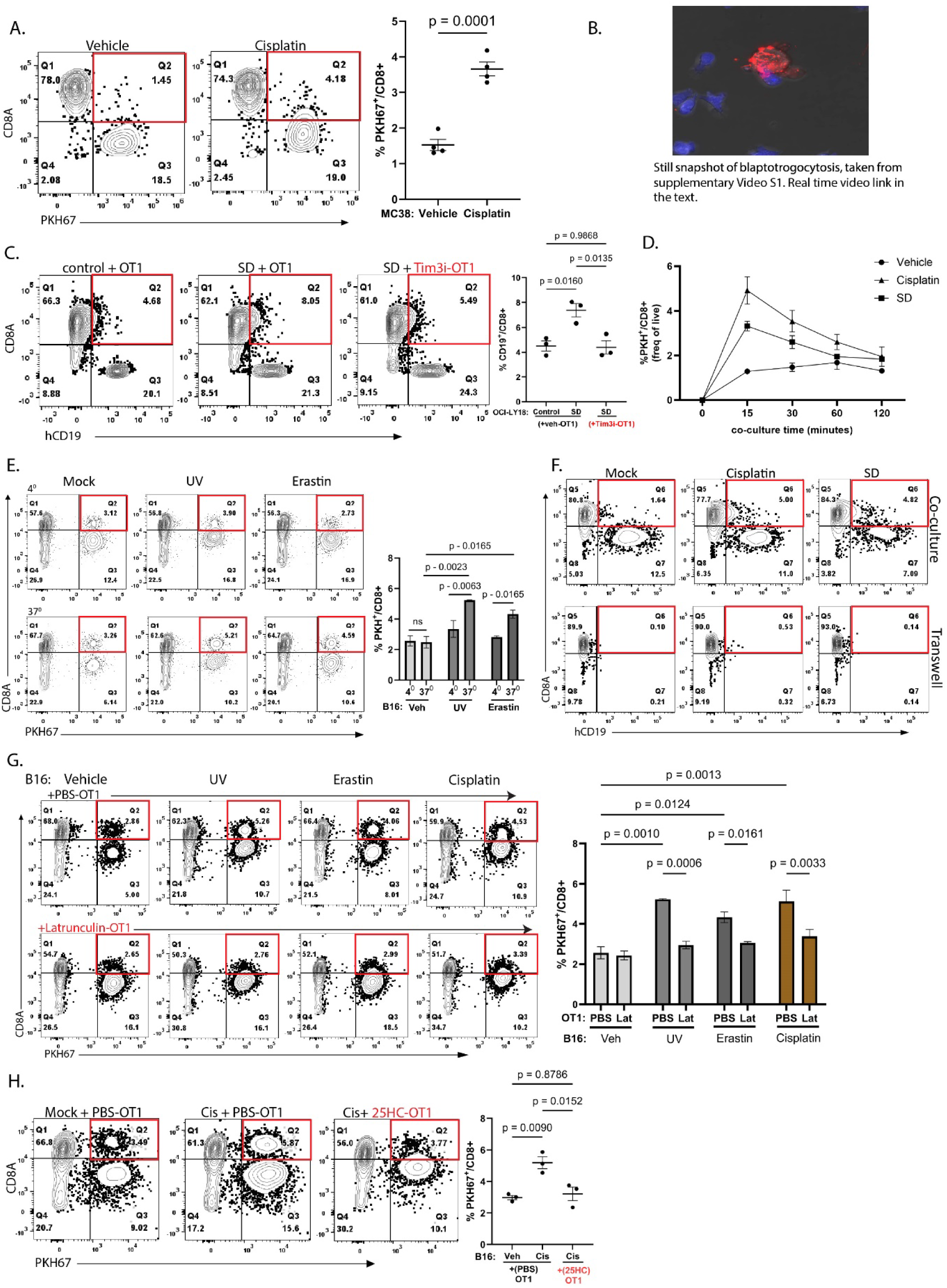
**a)** antigen-independent trogocytosis in the MC38 colon cancer model. Q2 - %PKH67+ CD8+ cells from vehicle-or cisplatin-treated MC38 + OT-1 co-cultures. **b)** Still frame from supplementary Video1 showing live cancer cell-CTL contact and membrane capture. Link for complete video: https://drive.google.com/file/d/1Yf-UrafFHT6NMPO17q0AIHoiQeqo_HXi/view?usp=sharing. **c)** Blaptotrogocytosis-mediated CD19 protein transfer from control or serum deprived (SD) OCI-LY18 cells onto OT-1 cells (%CD19+ CD8+). ***(d-h)*** characterization of blaptotrogocytosis: **d)** time course of membrane transfer from control, cisplatin-treated, or SD B16F10 cells onto OT-I CTLs over 15–120 min; **e)** temperature dependence of PKH67+ membrane transfer from damaged B16F10 cells onto OT-I CTLs at 4 °C versus 37 °C; **f)** contact dependence of CD19 transfer from control (mock), cisplatin-treated, or SD B16F10 cells onto OT-I CTLs cultured together (top) or separated by a Transwell membrane (bottom) for 15 min; **g)** actin dependence shown by PKH67+ membrane transfer from damaged cancer cells onto control or latrunculin-treated OT-I CTLs; and **h)** dependence on membrane fusion shown by PKH67+ membrane transfer from damaged-B16F10 cells onto control or 25-Hydroxycholesterol (25HC)-treated OT-1 CTLs. Data represent ≥3 independent experiments (mean ± SEM), unpaired two-tailed t-test **(*a*)**, one-way ANOVA (**c, h**), or two-way ANOVA (***e, g***).

**Supplementary Figure 2.**
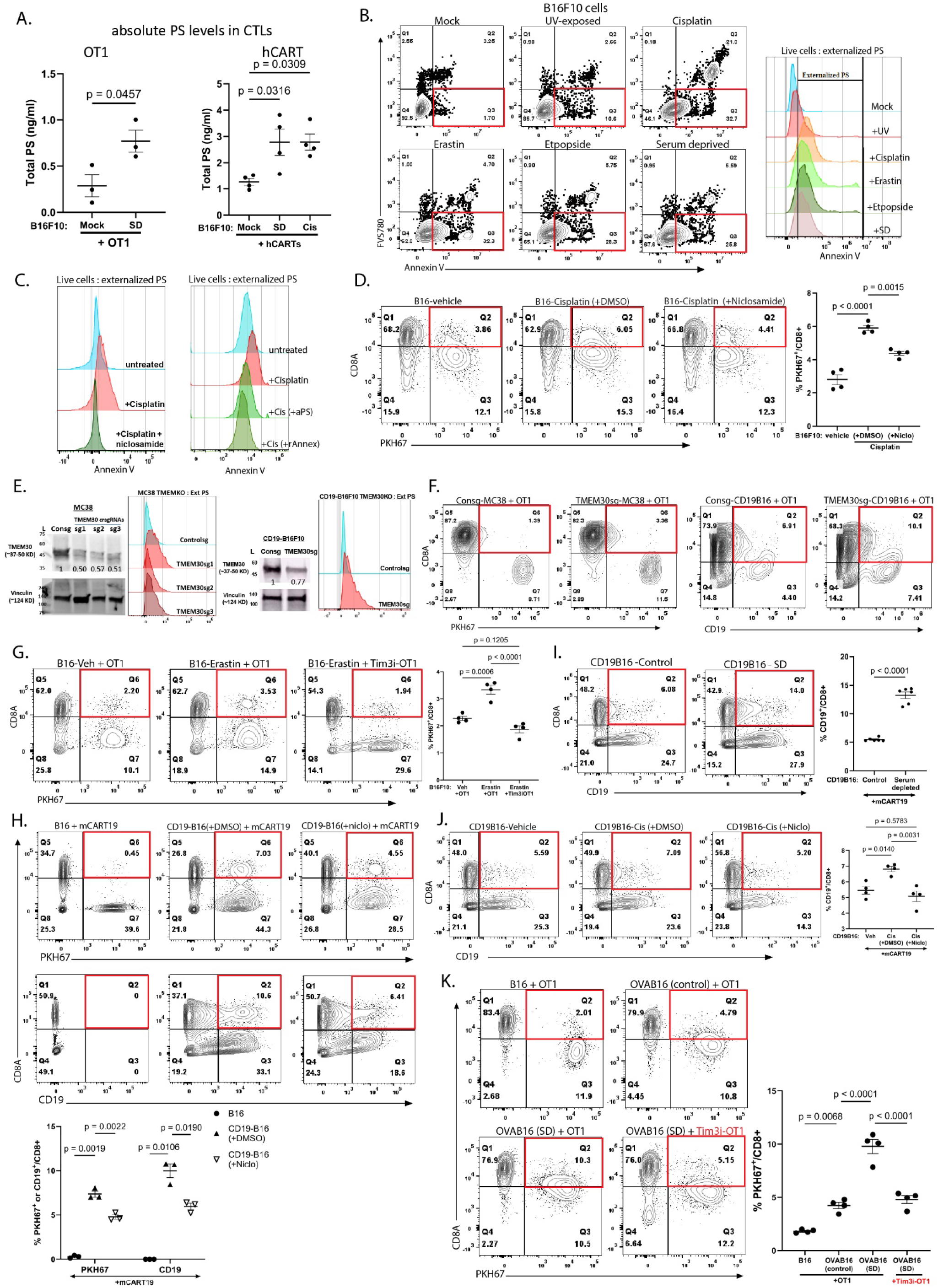
**a)** Total intracellular PS levels (ng/ml) in CTLs after co-culture with control or damaged B16F10 cells, measured by ELISA. **b)** surface PS expression on live B16F10 treated with UV irradiation, chemotherapy (cisplatin, erastin, or etoposide), or serum deprivation (contour plots showing AnnexinV+, FVS780-(Q3), and histograms show AnnexinV florescence on live CD8+ cells). **c)** surface PS blockage on cisplatin-treated cancer cells using ANO scramblase inhibitor niclosamide (left), or αPS or rAnnexin V (right). **d)** %PKH67+ CD8+ (of live cells) in (±niclosamide) cisplatin-treated B16F10 + OT-1 cocultures. **e)** Genetic model of constitutive PS exposure via *Tmem30a* flippase knockout in MC38 (left) and CD19-B16F10 (right) cells; immunoblots confirm *Tmem30a* loss and histograms show elevated surface PS. **f)** %PKH67+ CD8+ and %CD19+ CD8+ (of live cells) in control or *Tmem30a*-KO MC38 + OT-I or CD19-B16F10 + OT-I co-cultures. **g)** %PKH67+ CD8+ cells (of live) in erastin-treated B16F10 + control or Tim3i-treated OT-1 co-cultures. **h)** Antigen-driven trogocytosis model: % PKH67+ CD8+ (top) and % hCD19+ CD8+ cells (bottom) in ± niclosamide (PKH-labelled) hCD19-B16F10 + mCAR-T19 co-cultures. ***(i, j)*** %CD19+ CD8+ (of live cells) in mCART19 cocultures with - **i)** control or SD hCD19-B16F10, or **j)** control or ±niclosamide cisplatin-treated hCD19-B16F10. **k)** %PKH67+ CD8+ (of live cells) in parental, control- or SD-OVA-B16F10 cells co-cultured with untreated or TIM-3i–treated OT-1 CTLs. Data represent 3-6 experiments (mean ± SEM). Statistical significance was determined by unpaired two-tailed t-test ***(a(left), i)***, one-way ANOVA (***a(right), d, g, j, k***) or two-way ANOVA (***h***).

**Supplementary Figure 3.**
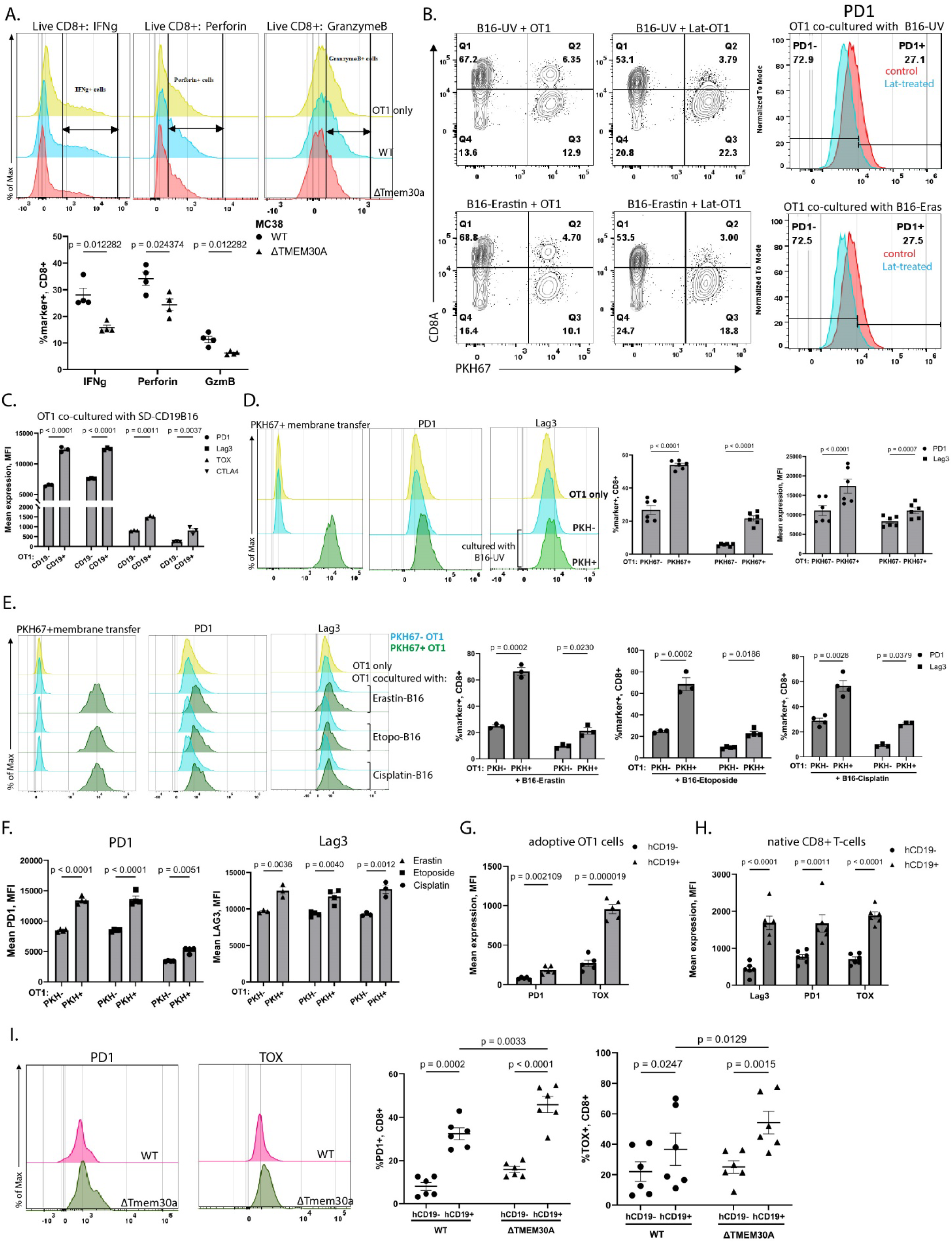
**a)** Offset histograms (normalized to mode) of IFNγ, perforin, and granzyme B in OT-I CTLs alone or co-cultured with control or *Tmem30a*-KO MC38 cells, quantified as % marker+ CD8+ cells (of live cells). **b)** Contour plots showing blaptotrogocytosis, and histograms show PD-1 expression (on live CD8+) in bulk control or latrunculin-treated OT-I CTLs co-cultured with damaged B16F10 cells. **c)** MFI of PD1, LAG3, TOX and CTLA4 for Fig. 3d). **d)** PD-1 and LAG-3 expression (% marker+ CD8+ quantification shown on the left; MFI on the right) on OT-1 CTLs alone or gated as TRG+ (PKH67+) or TRG-(PKH67-) OT-1 cells from co-cultures. **e)** PD-1 and LAG-3 histograms and % marker+ (within respective PKH67+ CD8+ or PKH67-CD8-gates) on OT1 CTLs alone or gated as TRG+ or TRG-from damaged-B16F10+OT-1 cocultures. **f)** MFI of *e)*. ***(g, h)*** MFI of markers on TRG+ (hCD19+) or TRG-(hCD19-) TILs from Fig. 3f) and Fig. 3g). **i)** PD-1 and TOX expression (%marker+ within live CD8+ hCD19+ or CD8+ hCD19-gates) on TRG+ CD8+ TILs from wild-type (WT) or *Tmem30a-KO* (ΔTMEM30) hCD19-B16F10 tumors in the adoptive transfer model (n=6 mice/group). Data represent 3-4 independent experiments (mean ± SEM); two-way ANOVA.

**Supplementary Figure 4.**
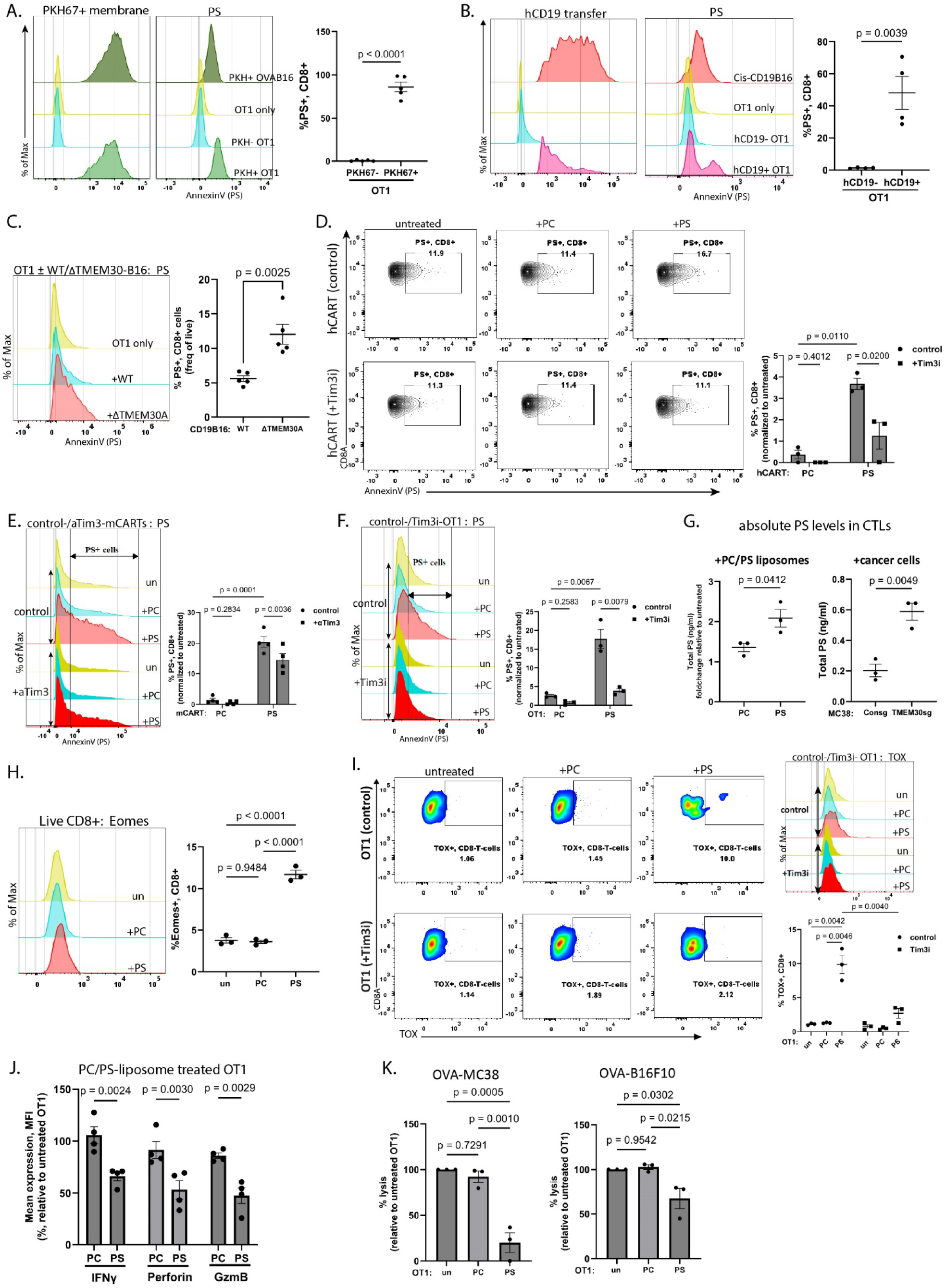
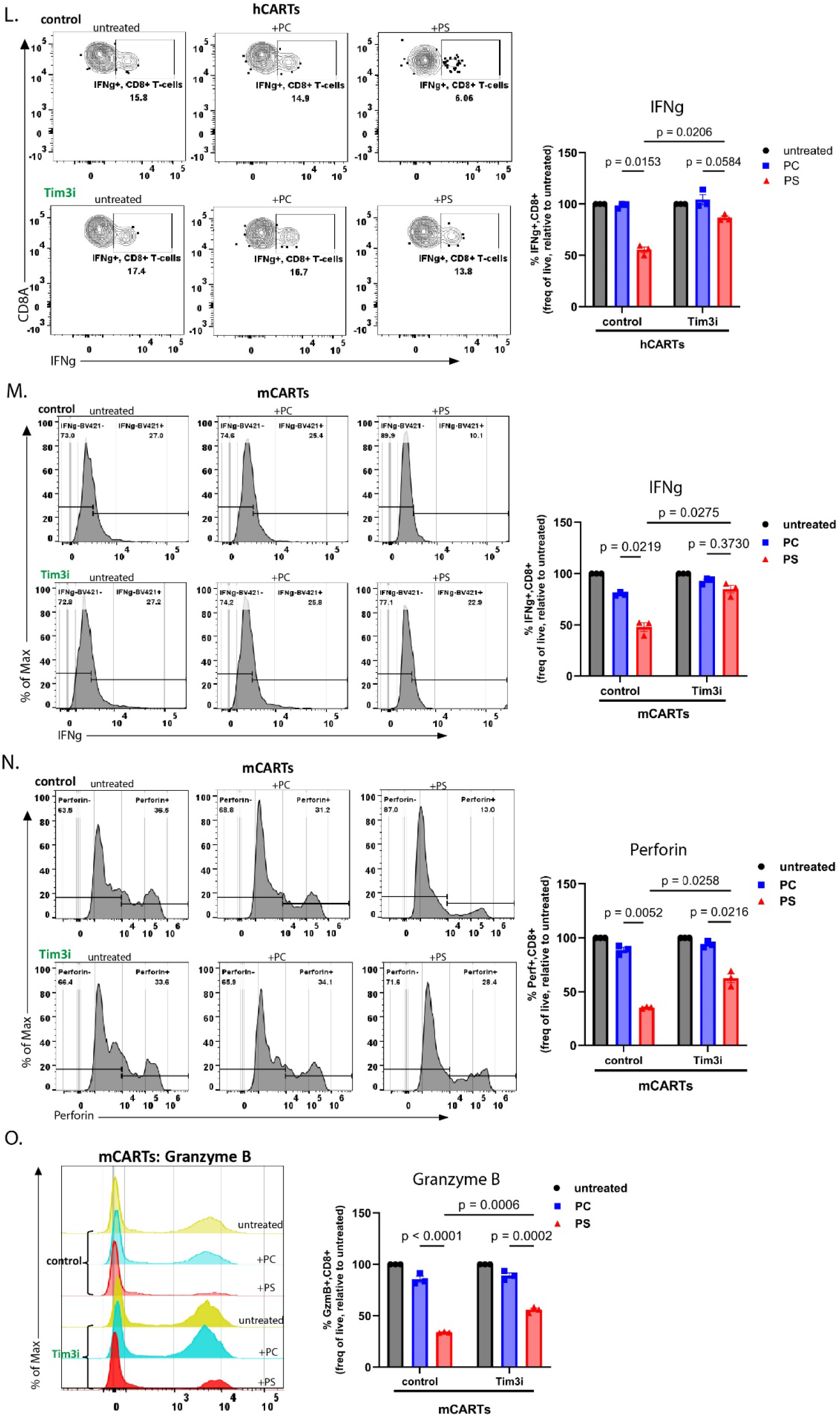
***(a–f)*** Surface PS expression (% PS+ CD8+) on: **a)** TRG+ (PKH67+) vs TRG-(PKH67-) OT-1 CTLs from OVA-B16F10+OT-1 co-cultures (of live PKH67+ or PKH67-CD8+ gates); **b)** hCD19+ vs. hCD19-OT-I CTLs from cisplatin-treated hCD19-B16F10 + OT-I co-cultures (within live hCD19+ or hCD19-CD8+ gates); **c)** OT-I CTLs co-cultured with control or *Tmem30a*-KO CD19-B16F10 cells (of live cells); **d)** PC/PS liposome-treated human aCD19-CARTs (±Tim3i); **e)** PC/PS liposome-treated mCARTs cells (±αTim3); and **f)** OT-I CTLs (± TIM-3i) (of live cells, normalized to respective untreated controls). **g)** Intracellular PS levels (ng/ml) in OT-I CTLs treated with PC/PS liposomes (left, fold change relative to untreated) or co-cultured with control or *Tmem30a*-KO MC38 cells (right). **h)** Eomes expression (% Eomes+ CD8+, of live CD8+ cells) in PC/PS-liposome-treated OT-I CTLs. **i)** contour plots and histograms (live CD8+) for TOX, and % TOX+ CD8+ cells (of live cells) in PC/PS-treated OT-I CTLs (±TIM-3i). **j)** MFI of IFNγ, perforin, and granzyme B in PC/PS-treated OT-I CTLs (relative to untreated) for Fig. 4f). **k)** Cytotoxic ability of PC/PS-treated OT-I CTLs towards OVA-MC38 or OVA-B16F10 cells (relative to untreated OT-I). ***(l–o)*** Effector molecule expression (%marker+ CD8+, of live cells, relative to respective untreated controls) in PC/PS-treated control or TIM-3i-pretreated: **l)** human αCD19 CAR-T cells (IFNγ), and **m–o)** mouse CAR-T19 cells – **m)** IFNγ; **n)** perforin; **o)** granzyme B. Data represent 3-5 independent experiments (mean ± SEM); unpaired two-tailed t-test (***a-c, g***), one-way ANOVA (***h, k***) or two-way ANOVA (***d-f, i-j, l-o***).

**Supplementary Figure 5.**
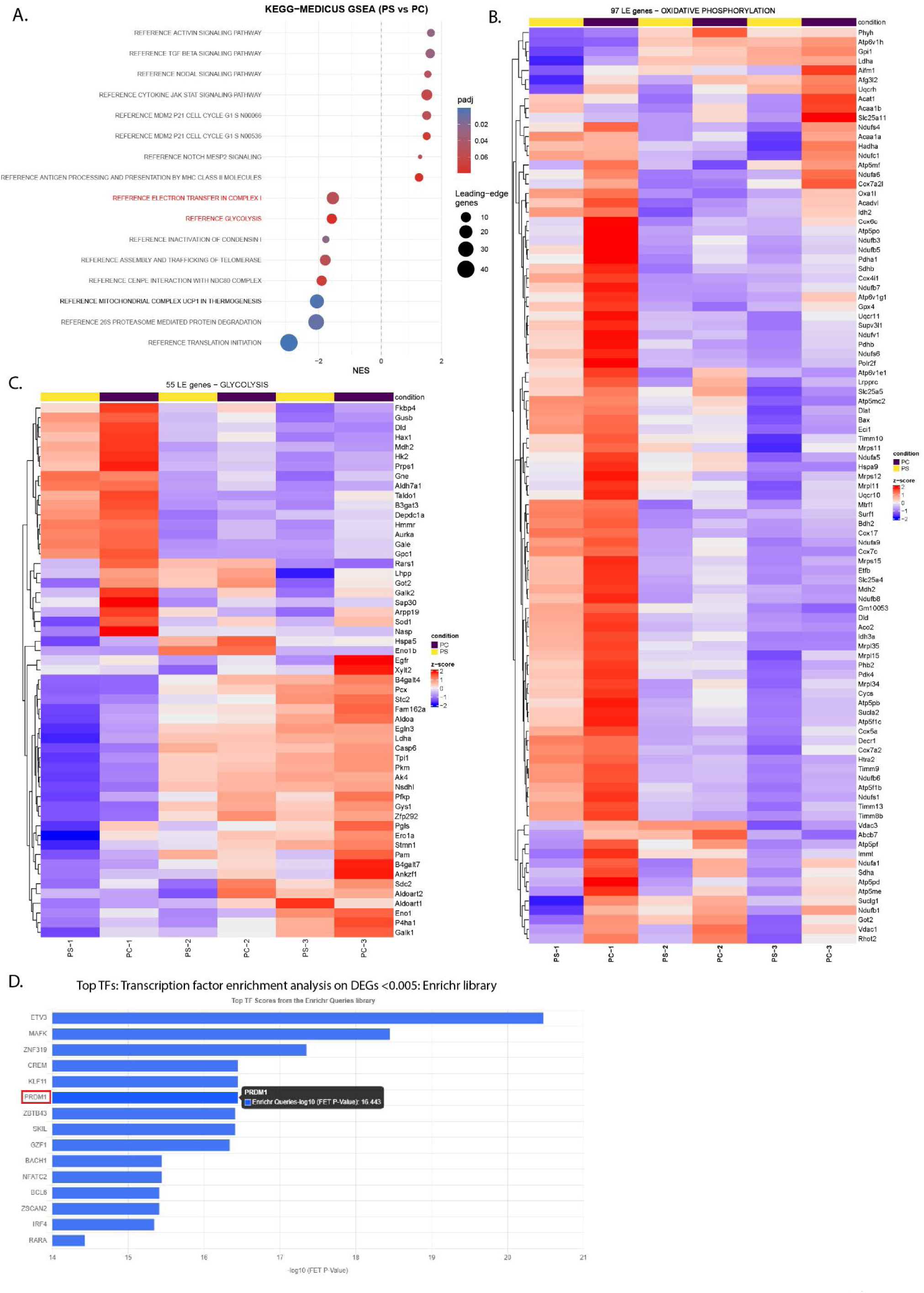
**a)** Bubble plot of significantly enriched KEGG Medicus pathways from RNA-seq data of PS-vs. PC-liposome-treated OT-1 CTLs. Notable molecular processes: glycolysis, mitochondrial respiration and electron transport chain are highlighted in red. ***(b, c)*** Heatmaps of leading-edge genes associated with **b)** oxidative phosphorylation and **c)** Glycolysis, in PS-treated OT-1 CTLs. **d)** Top 10 enriched transcription factors identified by ChEA3 (Enrichr library), ranked by −log10 Fisher’s Exact Test p-value.

**Supplementary Figure 6.**
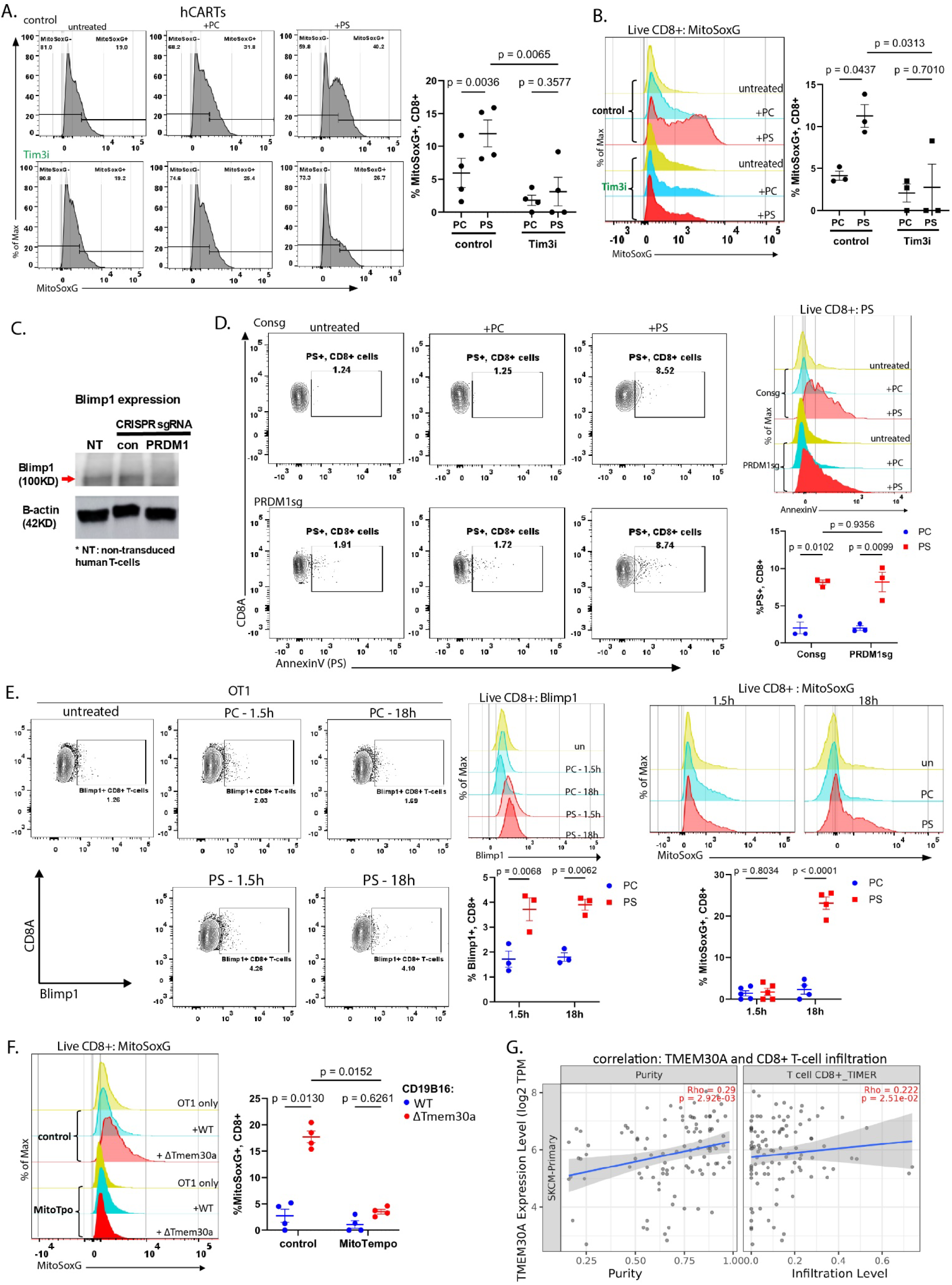
***(a, b)*** MitoSOX staining (%MitoSOX+ CD8+, of live cells, normalized to respective untreated controls) in PC/PS-liposome-treated control- or Tim3i-pretreated: **a)** hCART cells, and **b)** OT-1 CTLs. **c)** western blot confirming *PRDM1* knockout in human CAR-T cells transduced with control or *PRDM1* sgRNA (**ββ**-actin used as a loading control). **d)** surface PS expression (%PS+ CD8+) in PC/PS-liposome-treated control (Consg) or *PRDM1-KO* (*PRDM1*sg) human CD8+ T cells. **e)** Blimp-1 expression and MitoSOX fluorescence in OT-1 CTLs treated with PC or PS liposomes for 1.5 or 18 h. **f)** MitoSOX fluorescence (histograms, live CD8+) and %MitoSOX+ CD8+ cells (of live) in untreated or MitoTEMPO-pretreated OT-1 CTLs cocultured with WT or Tmeme30a-KO CD19-B16F10 cells. **g)** Purity-adjusted partial Spearman correlation between *TMEM30A* expression and CD8+ T-cell infiltration in SKCM (TIMER3; n = 470; ρ = 0.222, P = 0.025). Data represent 3-4 independent experiments (mean ± SEM). Statistical significance was determined by two-way ANOVA.

**Supplementary Figure 7.**
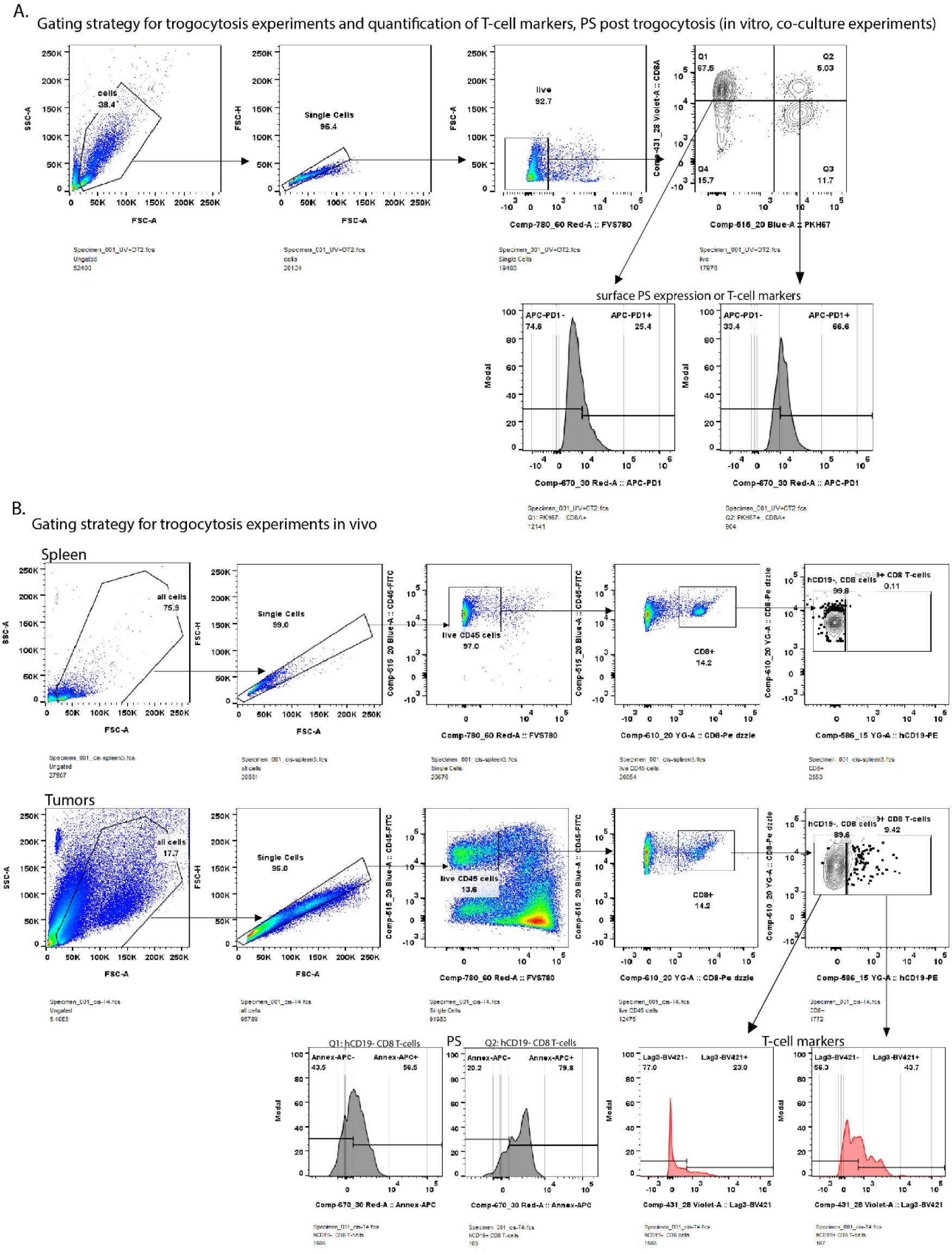
**a)** Gating strategy for quantifying *in vitro* trogocytosis and T-cell markers, effector molecules, and surface PS on TRG-(Q1) and TRG+ (Q2) CD8+ CTLs. **b)** gating strategy for quantifying trogocytosis and T-cell markers or surface PS on TRG- and TRG+ CD8+ CTLs isolated from tumors and spleens.

**Supplementary Figure 8.**
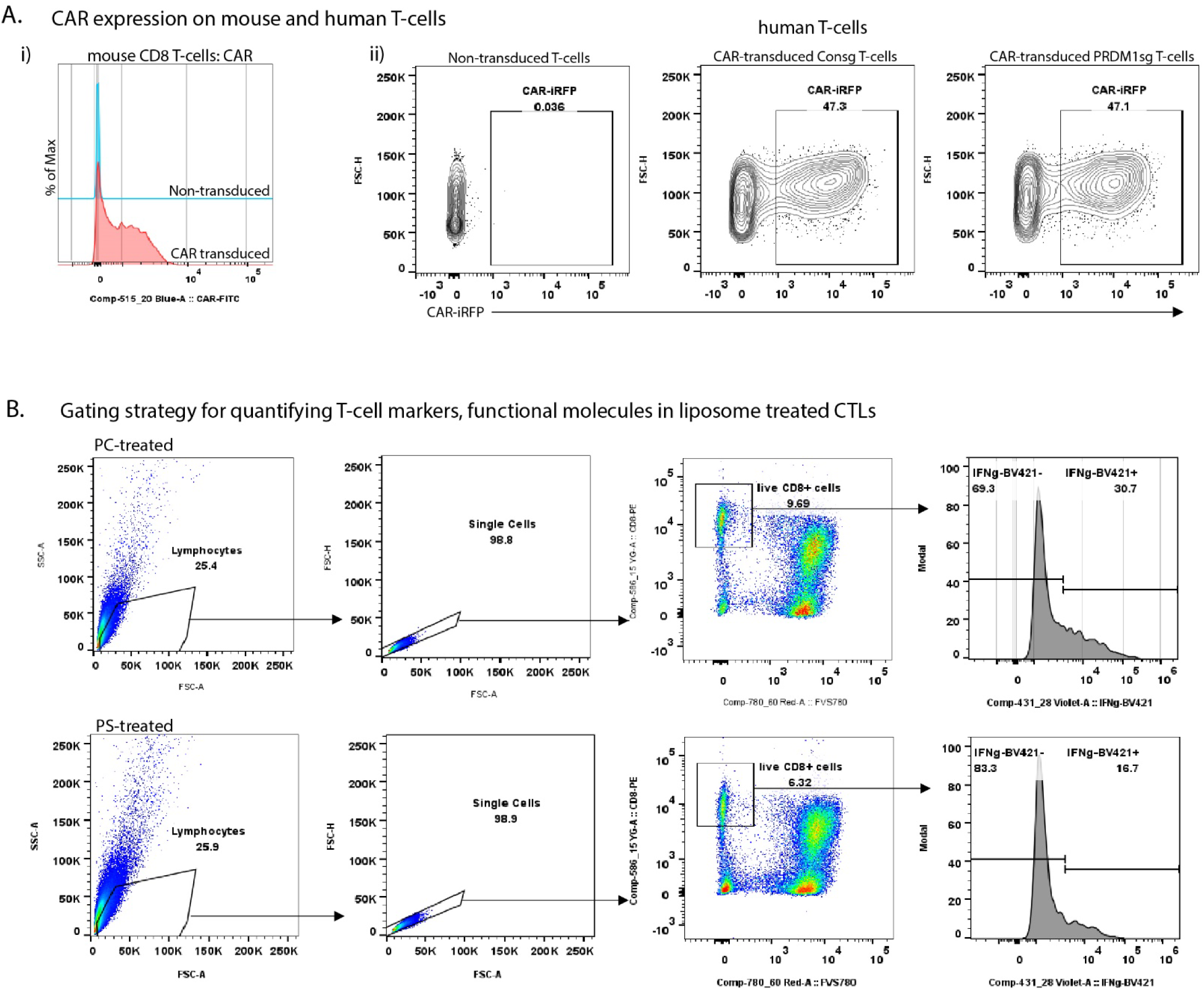
**a)** Confirmation of CAR expression on CAR-transduced - i) mouse CD8+ T-cells, and ii) human Controlsg (control) or *PRDM1*sg (Blimp1KO) T-cells. **b)** gating strategy for assessing the expression of T-cell state and functional markers in CD8+ T-cells following PC/PS-liposome treatment across all liposome experiments.

